# Single-cell long-read targeted sequencing reveals transcriptional variation in ovarian cancer

**DOI:** 10.1101/2023.07.17.549422

**Authors:** Ashley Byrne, Daniel Le, Kostianna Sereti, Hari Menon, Neha Patel, Jessica Lund, Ana Xavier-Magalhaes, Minyi Shi, Timothy Sterne-Weiler, Zora Modrusan, William Stephenson

## Abstract

Single-cell RNA sequencing predominantly employs short-read sequencing to characterize cell types, states and dynamics; however, it is inadequate for comprehensive characterization of RNA isoforms. Long-read sequencing technologies enable single-cell RNA isoform detection but are hampered by lower throughput and unintended sequencing of artifacts. Here we developed Single-cell Targeted Isoform Long-Read Sequencing (scTaILoR-seq), a hybridization capture method which targets over a thousand genes of interest, improving the median number of unique transcripts per cell by 29-fold. We used scTaILoR-seq to identify and quantify RNA isoforms from ovarian cancer cell lines and primary tumors, yielding 10,796 single-cell transcriptomes. Using long-read variant calling we revealed associations of expressed single nucleotide variants (SNVs) with alternative transcript structures. In addition, phasing of SNVs across transcripts facilitated measurement of allelic imbalance within distinct cell populations. Overall, scTaILoR-seq is a long-read targeted RNA sequencing method and analytical framework for exploring transcriptional variation at single-cell resolution.

## Introduction

Alternative RNA splicing is a key driver of proteome complexity and cellular phenotypic diversity. Approximately 95% of human multi-exon genes are alternatively spliced and 15-25% of human hereditary diseases and cancers are linked to alternative splicing^1–3^. Although short-read RNA sequencing has been widely adopted to measure gene expression, it remains challenging to identify full-length isoforms with only 20-40% of the human transcriptome being assembled using gold standard isoform reconstruction tools^4–6^. Additionally, alternative splicing, cleavage and polyadenylation events have been shown to be highly tissue-specific^7,8^. Thus, to better understand cellular diversity and dynamics in health and disease, high-resolution isoform-level transcriptomic information is required.

Single-cell RNA sequencing (scRNA-seq) has advanced our understanding of cellular heterogeneity, delivering transformative insights into a wide array of pathologies including autoimmune diseases^9,10^, neurological disorders^11,12^ and cancer^13,14^. To date, the vast majority of single-cell RNA profiling studies have employed short-read sequencing to measure gene expression which is typically quantified by counting reads derived from the 3’- or 5’-ends of genes. While useful for gene expression analysis, identification of isoforms remains challenging for single-cell short-read sequencing due to limited gene body coverage. To address this, multiple groups have performed long-read sequencing of cDNA from single cells which enables sequencing of full-length molecules^15–22^. However, to accurately identify cell barcodes (CBs) and unique molecular identifiers (UMIs), these strategies require short-read sequencing paired with long-read sequencing or specialized library preparation steps to improve read accuracy at the cost of sequencing throughput. Additionally, the majority of these studies have demonstrated sequencing of a relatively small number of cells at low per-cell sequencing depth due to current throughput limitations of long-read platforms^15,18,23,24^. Recent efforts have been developed to employ hybridization-based capture strategies to enrich selected genes of interest^17,25^. Gene panel designs utilized in previous studies have typically focused on specific biological questions and encompassed less than 50 target genes, which presents a challenge for cell annotation in complex tissues and requires additional short-read sequencing^17,25^. A particular issue inherent to single-cell long-read library preparation and sequencing is the presence of unwanted artifacts that consume valuable sequencing throughput. These artifactual reads do not exhibit the expected cDNA structure after reverse transcription and amplification; rather, often contain template switching by-products or lack adapter sequences^16^.

To address the aforementioned shortcomings, we have developed single-cell targeted isoform long-read sequencing (scTaILoR-seq). scTaILoR-seq makes use of commercially-available or custom-designed gene panels to enrich for greater than 1,000 genes of interest. Additionally, scTaILoR-seq mitigates the presence of artifacts common to single-cell RNA-seq cDNA by enriching for molecules with the expected adapter sequence using biotinylated PCR primers. Following both gene panel enrichment and artifact mitigation, nanopore long-read sequencing is used to enable assignment, identification and quantification of transcript isoforms in thousands of single cells.

Using scTaILoR-seq, we characterized differential transcript expression among three ovarian cancer cell lines. To identify CBs and UMIs, short-read guided (*SiCeLoRe)* and unguided (*wf-single-cell)* assignment approaches were compared, resulting in high concordance. We then applied scTaILoR-seq with the unguided CB/UMI assignment method *wf-single-cell* (*i.e.* sans supplemental short-read sequencing) to profile dissociated tumor cells (DTCs) from two ovarian cancer patients. This enabled identification of isoforms, reconstruction of immune repertoires and detection of expressed single nucleotide variants (SNVs) at the single-cell level. Additionally, long-reads enabled SNV phasing to assemble haplotypes and estimate allelic imbalance from individual tumor epithelial cells. Collectively, scTaILoR-seq establishes an efficient approach for sensitive characterization of diverse transcript variants in single cells.

## Results

### Targeted gene enrichment method development

To evaluate gene enrichment, we performed droplet-based single-cell 3’-end RNA sequencing on a mixture of three human ovarian cancer lines (SK-OV-3, COV504, and IGROV-1) (**Fig. 1**). The single-cell cDNA from this cell line mixture was used to evaluate a variety of strategies aimed at determining optimal conditions for long-read sequencing (**Fig. 2a, Supplementary Fig. 1**). Preliminary assessment of the targeted approach using a 10x Genomics pan-cancer probe panel demonstrated efficient enrichment of cancer-associated genes with short-read sequencing (**Supplementary Notes**). We next sought to maximize the proportion of complete reads (*i.e.* reads containing both the template switch oligo (TSO) adapter and poly(A) sequences) using long-read sequencing. A previously described artifact mitigation (AM) approach was deployed to reduce TSO-TSO byproducts from library preparation^16^ using biotinylated PCR primers complementary to the Read1 sequence, which enabled streptavidin-coated magnetic bead pull-down and subsequent amplification of complete cDNA constructs. Compared to the targeted approach without AM, the targeted+AM strategy displayed an 11.8% increase in complete read proportion concomitant with a marked decrease in TSO-TSO artifacts (**Fig. 2b**). Next, we investigated an orthogonal TSO-TSO depletion approach based on circularization of targeted complete cDNA using rolling circle amplification to concatemeric consensus (targeted+R2C2)^16,19^(**Methods**). Compared to targeted+AM, the targeted+R2C2 approach exhibited a slightly higher proportion of complete reads and fewer TSO-TSO artifacts; however, it yielded much lower read throughput: 4.4M versus 18M average passed reads per flow cell compared to the targeted+AM approach (**Fig. 2b**). Therefore, the targeted+AM strategy displayed an optimal balance between increased complete read proportion and higher read throughput. For this reason, the targeted+AM approach became the basis of scTaILoR-seq, which was employed in subsequent targeted experiments.

**Fig. 1.**
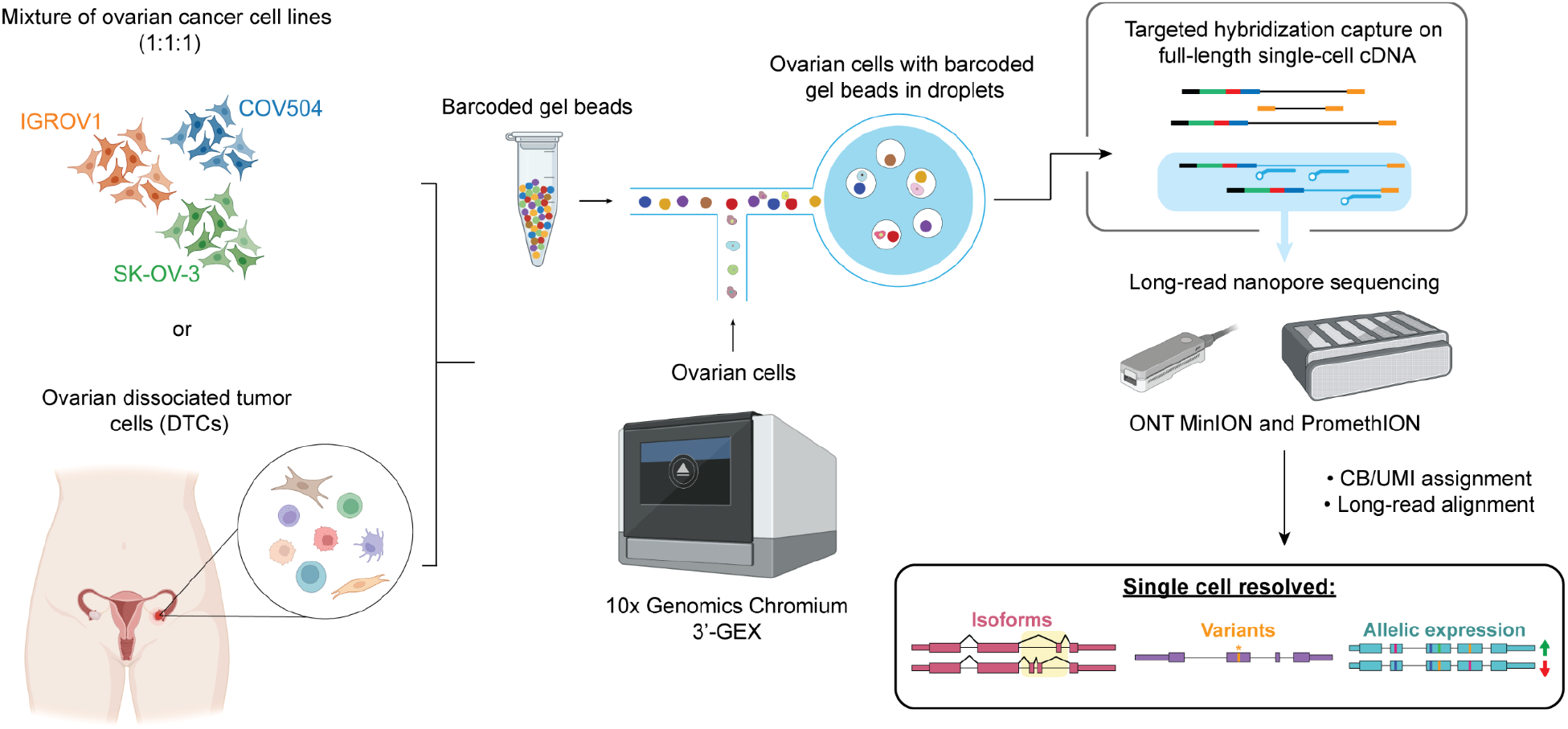
Overview of targeted single-cell long-read sequencing. Ovarian cell lines or dissociated tumor cells are processed using droplet-based single-cell RNA-seq 3’-Gene expression assay to obtain cDNA. Targeted enrichment is performed followed by nanopore sequencing, cell barcode (CB) and unique molecular identifier (UMI) assignment and long-read alignment. Downstream analysis enables measurement of isoforms, SNVs and allelic expression at single-cell resolution.

**Fig. 2.**
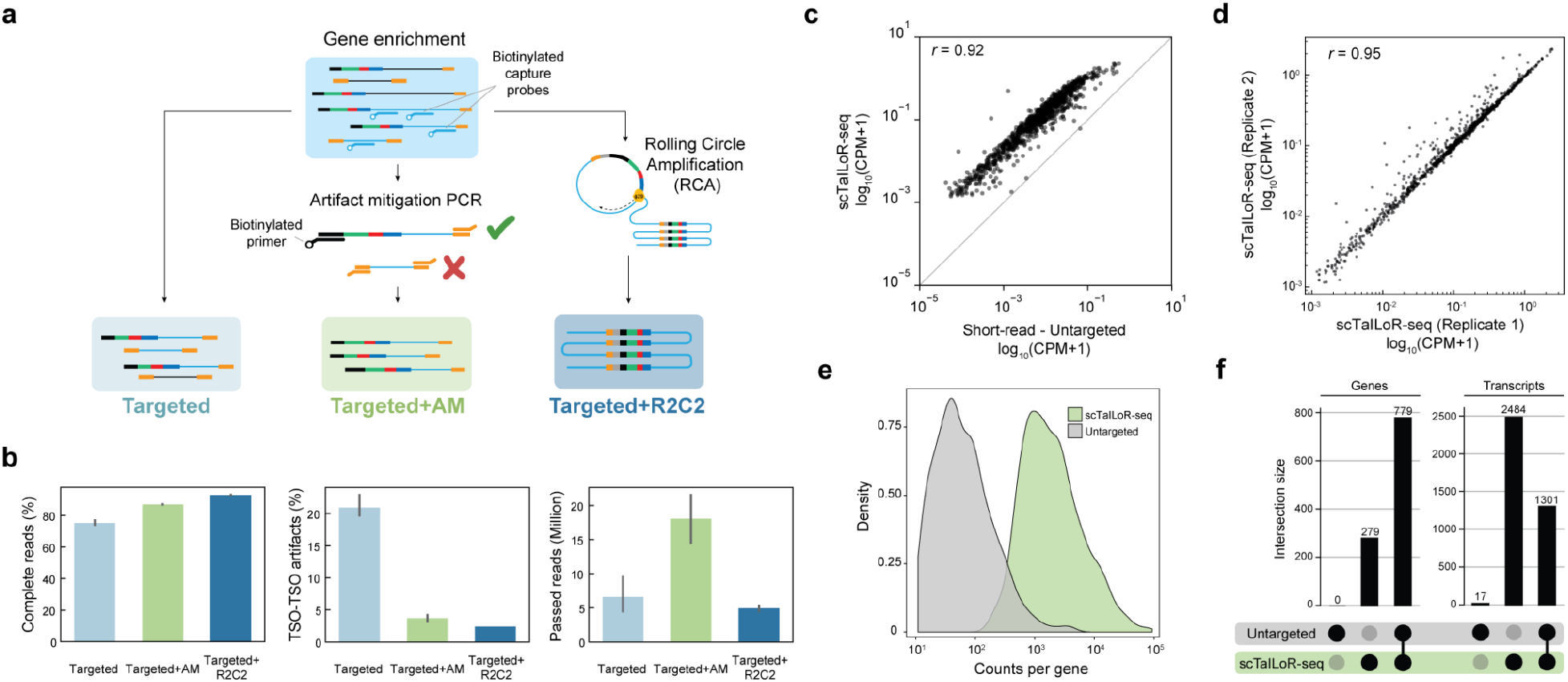
Targeted long-read sequencing optimization. **a**, Schematic detailing library preparation methods (targeted, targeted+AM and targeted+R2C2) tested for enrichment based long-read sequencing. **b**, Complete reads (left), TSO-TSO artifacts (middle) and number of passed reads (right) across library preparation methods. **c**, Pseudobulk gene-level expression between short-read untargeted and scTaILoR-seq library preparation methods. Log10(CPM +1), CPM - Counts per million. **d**, Pseudobulk gene expression correlation for scTaILoR-seq across replicates. **e**, Counts per gene density distributions for untargeted and scTaILoR-seq. **f**, Number of genes and transcripts uniquely detected (single dot) or shared (‘connected’ or ‘joint’ dots) across untargeted and scTaILoR-seq.

Next, we investigated the efficiency and reproducibility of scTaILoR-seq by measuring on-target gene expression levels. From read-depth normalized samples, we observe highly correlated mean gene expression (*r*=0.92) between scTaILoR-seq and untargeted (*i.e.* no gene enrichment) short-read sequencing, indicating that scTaILoR-seq faithfully recapitulates quantitative expression patterns (**Fig. 2c and Supplementary Fig. 2**). Gene expression was strongly correlated across replicates (*r*=0.95) with 98.8% gene overlap demonstrating the reproducibility of scTaILoR-seq (**Fig. 2d**). Moreover, scTaILoR-seq resulted in a 16.4-fold increase of on-target reads compared to untargeted long-read sequencing, yielding a significant boost in read counts per gene (Mann-Whitney *U* test, *P*=3.7×10^-129^) (**Fig. 2e**). Finally, scTaILoR-seq identified an additional 279 on-target genes and 2,484 annotated transcripts that were not detected in the long-read untargeted approach, showcasing the increased sensitivity enabled by enrichment (**Fig. 2f**). These results demonstrate that scTaILoR-seq improves gene and transcript detection via efficient allocation of sequencing reads to targeted genes, while preserving quantitative information.

Because higher error rates observed in nanopore sequencing reads can confound CB and UMI assignment, strategies that leverage supplemental short-read sequencing data to guide assignments have been developed^16,24^. We compared one such guided method, *SiCeLoRe*^16^, with an unguided approach, *wf-single-cell* (**Methods**). This unguided method eliminates the requirement for supplemental short-read sequencing to assign CBs and UMIs (**Supplementary Fig. 3a**). We observed a high degree of overlapping CBs between *SiCeLoRe* and *wf-single-cell* assignments. These overlapping CBs encompass nearly all of those found in the associated untargeted short-read sequencing data (**Supplementary Fig. 3b**). In addition, UMI counts per CB from *SiCeLoRe* and *wf-single-cell* were highly correlated (*r*=0.97) (**Supplementary Fig. 3c**). Similarly, gene expression for matched cell line populations was also highly correlated (**Supplementary Fig. 3d**). Taken together, these results indicate that scTaILoR-seq is compatible with current guided and unguided CB/UMI assignment methods.

### scTaILoR-seq enables detection of alternative splicing at the single-cell level

We sought to quantify the enrichment performance of scTaILoR-seq at the single-cell level using genetically-deconvoluted cell populations from the ovarian cell line mixture (**Fig. 3a**). Relative to the untargeted approach, scTaILoR-seq exhibited a 10-fold median increase in on-target genes per cell and a 29-fold median increase in both on-target UMIs and transcripts per cell (**Fig. 3b**). Additionally, the top-25 expressed genes from scTaILoR-seq were noticeably depleted of mitochondrial and house-keeping genes that were abundant in the untargeted approach (**Supplementary Fig. 4**). Next, we assessed whether scTaILoR-seq can be used to identify alternative splicing events across the ovarian cancer cell lines. Using differential transcript expression (Welch’s *t*-test), we identified significant cell line-specific isoform usage (Benjamini-Hochberg adjusted *P*<0.05; Methods) (**Supplementary Fig. 5**). For example, we identified alternative 5’ splice site usage of exon 2 in PARP2, the frequency of which varied across the three cell lines (**Fig. 3c and Supplementary Fig. 6**). Exon 2 of PARP2 is localized at the N-terminal region which is known to facilitate activation on DNA single strand breaks. Alternative splicing within this region may modulate the DNA damage sensing activity of PARP2^26^. Additionally, we identified a predominant alternative 5’-UTR and first exon usage event in the Rho-binding domain of RTKN specific to SK-OV-3 (**Fig. 3d and Supplementary Fig. 7**). RTKN is a scaffold protein that interacts with GTP-bound Rho proteins to subsequently regulate cell growth and transformation^27^. Overall, we demonstrated the ability of scTaILoR-seq to enrich for genes of interest, which enabled identification of differential isoform usage events and alternative splicing patterns at the single-cell level.

**Fig. 3.**
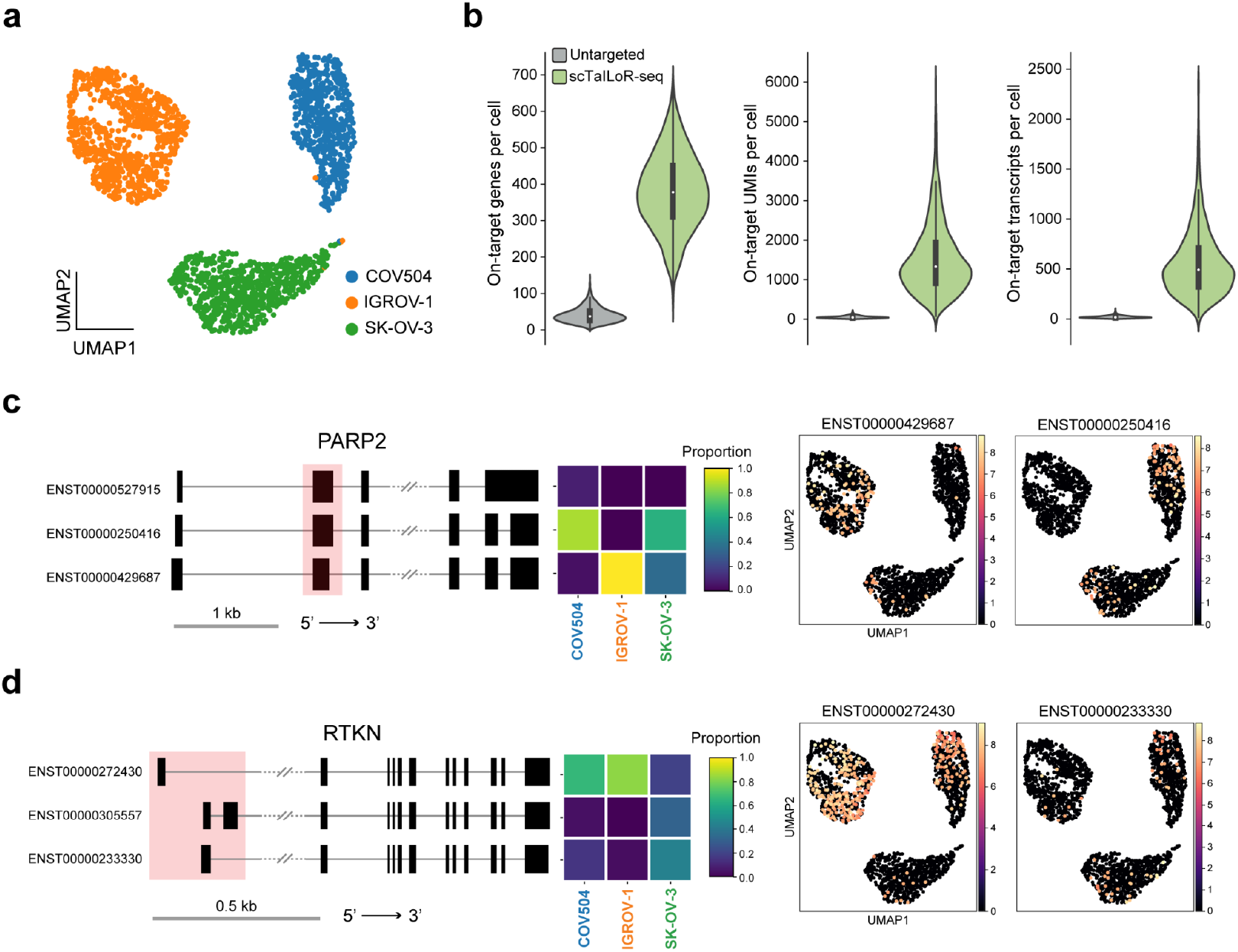
Single-cell enrichment metrics and cell-line specific alternative splicing. **a**, UMAP visualization of 3 mixed (1:1:1) ovarian cell lines (COV504, IGROV-1, and SK-OV-3). **b**, Comparison of on-target genes, UMIs and transcripts per cell across untargeted and scTaILoR-seq library preparation methods. **c**, Cluster level PARP2 isoform proportions and single-cell transcript UMAP visualization. Alternative 5’ splice site within exon 2 of PARP2 is indicated by the shaded pink rectangle in the transcript model. **d**, Cluster level RTKN isoform proportions and single-cell transcript UMAP visualization Alternative 5’-UTR and first exon usage of RTKN is indicated by the shaded pink rectangle in the transcript model.

### Surveying the transcriptional landscape of an ovarian tumor microenvironment with scTaILoR-seq

The tumor microenvironment (TME) is a complex niche characterized by dynamic interactions among diverse cell types including epithelial, stromal and immune cells. To quantify differential isoform usage and to annotate cell type populations within the TME, both pan-cancer and immune enrichment panels were used to target a total of 2,243 genes. We performed scTaILoR-seq on dissociated tumor cells (DTCs) from two stage-III treatment-naive ovarian cancer patients: P1 - high grade serous ovarian carcinoma (HGSOC), P2 - ovarian clear cell carcinoma (OCCC). Long-read sequencing was performed on the PromethION instrument resulting in a total of 371 million reads with a median of 4,020 and 1,867 UMIs per cell for P1 and P2, respectively. We detected 8,695 cells derived from the two patient samples and identified 5 major cell groups (B cells, T/NK cells, myeloid, fibroblast and epithelial) (**Supplementary Fig. 8**). Lineage-specific cell proportions were consistent between scTaILoR-seq and untargeted short-read data generated from the same single-cell cDNA (**Supplementary Fig. 9**). Of particular interest was sample P1 (*n*=2,482 cells) which was analyzed to a greater extent since it contains a higher number of EPCAM+ tumor epithelial cells (*n*=1,498) in addition to an even representation of both stromal and immune cells (**Fig. 4a**). Differential expression analysis identified transcripts from genes that were consistent with annotated cell identity such as expression of EPCAM in epithelial cells, multiple isoforms of COL3A1 and COL1A2 in fibroblasts, CD3E and CD2 in T cells and distinct C1QB isoforms in cells derived from the myeloid lineage (**Fig. 4b**).

**Fig. 4.**
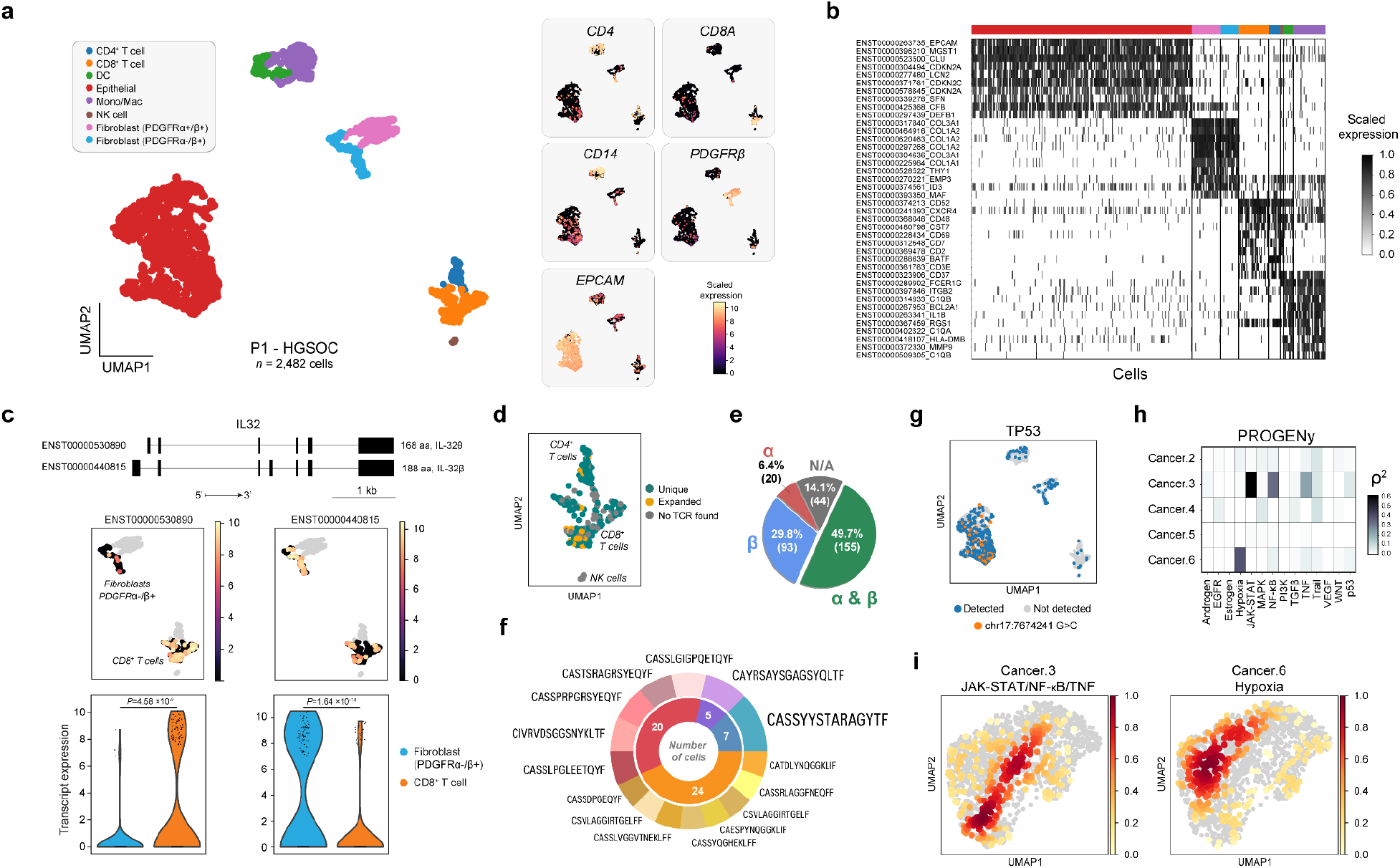
Profiling ovarian tumor cells with scTaILoR-seq. **a**, UMAP visualization of dissociated tumor cells (n=2,482) from a HGSOC patient (P1). Scaled expression of cell-type specific canonical marker genes are shown as additional UMAPs (right). **b**, Scaled expression of the top-10 differentially expressed transcripts across single cells. The colored horizontal bar at top corresponds to cell type annotations in **a**. **c**, IL-32 transcript models for theta and beta isotypes and differential IL-32 isoform usage identified across CD8+ T cells and PDGFRa-/b+ Fibroblasts (Mann-Whitney *U* test). **d**, Zoomed in UMAP of T cells showing successfully reconstructed TCRs. **e**, Proportion of T cells with TCR chain assignments: no chain identified (N/A), one chain identified (α or β) or both chains identified (α & β). **f**, Higher-order (>2 cells) clonotypes identified within T cells. The inner ring denotes the number of cells while the outer ring denotes individual clonotoype frequency. **g**, Projection of TP53 mutation (chr17:7674241_G>C) identified using *Clair3* on UMAP of dissociated tumor cells. **h**, Correlation between single-cell MSK HGSOC geneset expression and PROGENy pathway activity scores. **i**, MSK HGSOC Cancer.3 (JAK-STAT/NF-kB/TNF-active) and Cancer.6 (Hypoxia-active) geneset scores mapped to epithelial cell embedding.

Provided that alternative splicing events are prevalent in cancer and the associated TME^28,29^, we analyzed differential isoform usage between all cell types. This analysis identified 43 significant events (**Supplementary Table 2**) including differential IL-32 isoform usage between CD8+ T cells and PDGFRɑ-/β+ fibroblasts (Mann-Whitney *U* test; ENST00000530890 *P*=4.6×10^-9^ and ENST00000440815; *P*=1.6×10^-14^) (**Fig. 4c**). Expression of IL-32β isotype (ENST00000440815) was dominant across all cell types; whereas, IL-32θ (ENST00000530890) expression was markedly low in PDGFRɑ-/β+ fibroblasts (**Supplementary Fig. 10**). IL-32β is associated with hypoxic conditions in solid tumors and IL-32θ inhibits NF-kB which counters the epithelial-mesenchymal transition^30^.

Next, we turned our attention to the immune component of the TME, where current single-cell TCR/BCR reconstruction with short-read sequencing requires supplemental library preparation and is limited to 5’-expression profiling. A recent single-cell long-read study was unable to obtain sufficient read depth for low abundance TCR transcripts, indicating a need for increased detection sensitivity^31^. Thus, we asked whether scTaILoR-seq (3’-expression) would be amenable to TCR repertoire profiling. scTaILoR-seq reads were processed by *TRUST4* which performs single-cell repertoire reconstruction of TCR sequences^32^. Of the barcodes associated with successfully assembled TCRs, 98% corresponded to annotated T cells (**Fig. 4d**) and 85.9% had at least one chain (α and/or β) identified (**Fig. 4e**). With scTaILoR-seq, we obtained a TCR α/β chain pairing rate of 49.7%, which is a two-fold improvement over previous targeted and untargeted long-read strategies^17,33^. Within the expanded T cell population (*n*=56 cells), we identified 15 high-order clonotypes with more than two cells sharing identical CDR3 regions. The CDR3 sequence CASSYYSTARAGYTF was detected in seven cells, representing the largest observed clonotype population (**Fig. 4f**).

Characterization of the epithelial cell population was of particular interest because the DTCs were derived from epithelial ovarian tumors. Long-read sequencing enables detection of expressed SNVs that are outside the typical read length of short-read single-cell sequencing. For example, using the long-read variant caller *Clair3*^34^, we detected a SNV in exon 7 of the tumor suppressor TP53 (chr17:7674241 G>C). This missense variant (S241C) alters the DNA-binding domain of TP53 and is a likely pathogenic ovarian cancer mutation^35^. This SNV was detected exclusively in a sub-population of epithelial cells, despite TP53 expression across several other cell types (**Fig. 4g**). To further assess cancer-associated expression patterns among the epithelial cells, we performed pathway activity analysis using *PROGENy* which identified two signatures: JAK-STAT/NF-κB/TNFɑ and hypoxia (**Fig. 4h**). These two pathway activities were correlated with gene expression patterns characteristic of tumor cells from treatment-naive HGSOC patients: Cancer.3 and Cancer.6, respectively^13^. *PROGENy* signatures and associated gene expression patterns were localized to distinct cell subsets within the epithelial cell embedding (**Fig. 4i and Supplementary Fig. 11a-d**). Collectively, these data suggest that a large fraction of the epithelial cells exhibit distinguishing cancer signaling pathways consistent with ovarian cancer.

Overall, scTaILoR-seq effectively profiled the TME, enabling the discovery of cell type-specific isoforms, high-sensitivity TCR reconstruction, identification of expressed SNVs and detection of cancer-associated gene signatures.

### Identifying structural transcript variation associated with expressed SNVs

After determining the ability of scTaILoR-seq to detect SNVs, we asked whether these expressed variants were associated with differential transcript structures in HGSOC, as reported in several other tissues and cell lines^36^. We utilized a deep learning-based model called *SpliceAI*^37^ to predict and score cryptic splicing events associated with detected SNVs within the tumor epithelial cell population (**Fig. 5a**). For the 82 hits from 1,669 SNVs (*SpliceAI* score>0.1) (**Fig. 5b**), we identified transcript structure variation by assessing the coverage divergence (1-*r*^2^) between reads matching the reference base (REF) or the alternative base (ALT) of a given SNV site (**Fig. 5a**). Among the 82 queried hits, 44 displayed non-zero coverage divergence, indicating a difference of transcript structure between REF and ALT alleles. The local extent of coverage divergence was used to classify transcript structural events into two categories: “CDS” for protein-coding regions and “UTR/Intron” for untranslated regions and introns (**Fig. 5c**). We detected differential ELF3 transcript structures associated with chr1:202011127 A>C in exon 2 for which flanking retained introns were observed among REF reads (1,130 UMIs in 574 cells); whereas ALT reads (2,962 UMIs in 1,013 cells) exhibited normal splicing (**Fig. 5d**). ELF3 is a transcription factor strongly expressed in epithelial tissue and has been shown to inhibit the epithelial-to-mesenchymal transition^13,38^ while supporting angiogenesis^39^. Another example of differential transcript structures linked to an SNV (chr2:190975811 C>A) was observed with the transcription factor STAT1, which exhibited distinct allele-specific events (REF = 2,501 UMIs in 525 cells and ALT = 1,254 UMIs in 353 cells) that spanned both CDS and UTR/Intron (**Fig. 5e**). Taken together, scTaILoR-seq can provide insight into variation of transcript structures associated with SNVs, leading towards enhanced understanding of transcriptional complexity associated with genetic alteration in cancerous cells.

**Fig. 5.**
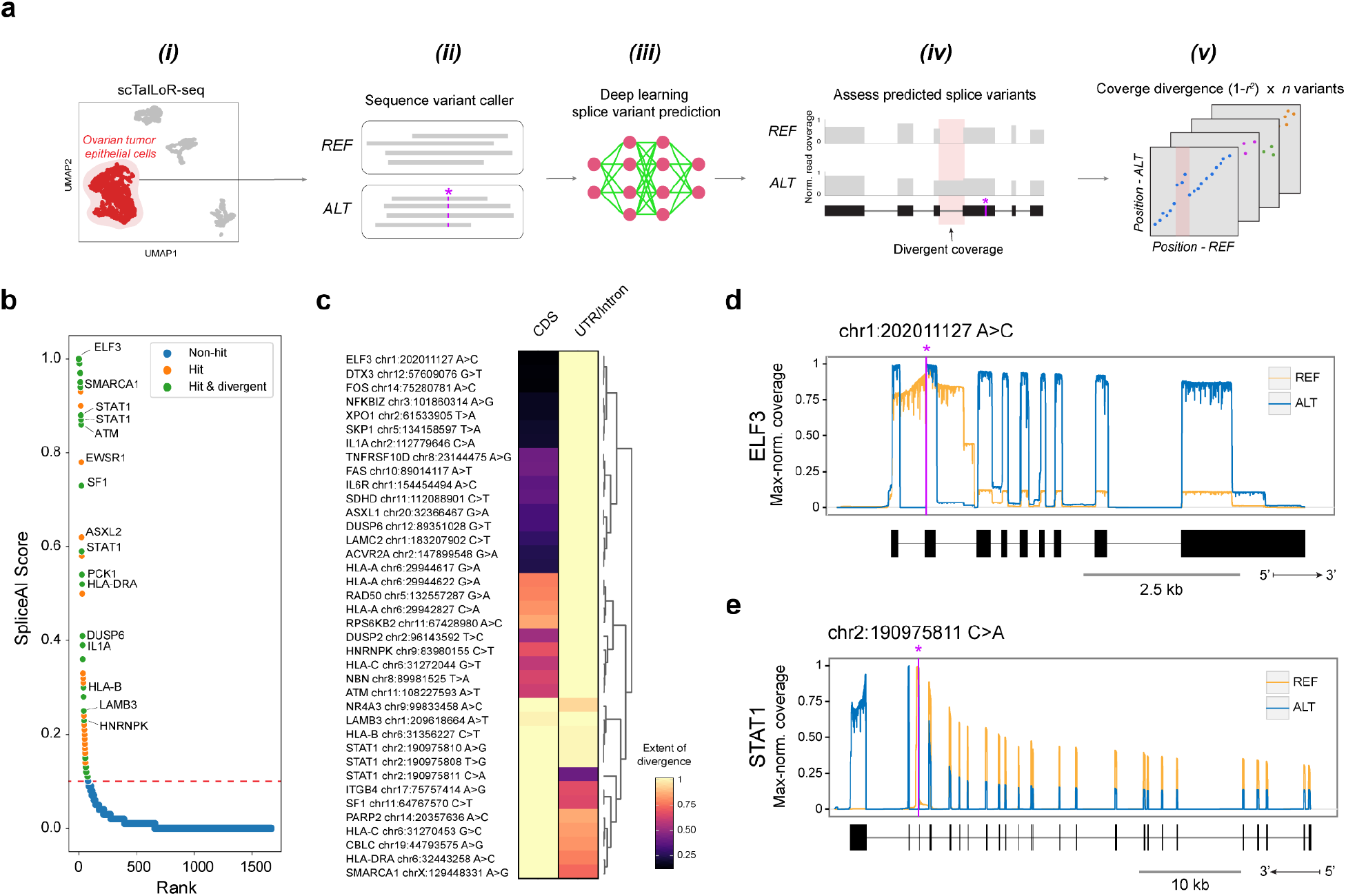
Identification of SNV associated differential transcript structures. **a**, Workflow for detecting SNV associated differential transcript structures: (i) Obtain reads from ovarian tumor epithelial cells (scTaILoR-seq), (ii) Determine SNVs using *Clair3*, (iii) Predict cryptic splice events using *SpliceAI*, (iv) Compute coverage divergence between reference (REF) or alternative (ALT) variant reads, (v) Identify differential transcript structures among *SpliceAI* hits using coverage divergence. **b**, SNVs of genes exhibiting *SpliceAI* score above threshold value of 0.1. Each SNV is colored by *SpliceAI* score (“Non-hits” - blue, “Hits” - orange) and whether a hit also displays coverage divergence between REF and ALT (“Hit and divergent” - green). **c**, Hierarchical clustering based on extent of divergence at transcript structural elements (CDS and UTR/Intron) of 44 “Hit and divergent” SNVs. **d** and **e**, Plots for ELF3 and STAT1, respectively: normalized coverage tracks for REF and ALT and corresponding transcript model.

### Phasing of expressed SNVs reveals allelic imbalance within tumor epithelial cells

Given the long-read output of scTaILoR-seq, we reasoned that transcripts containing multiple SNVs could be used for haplotype reconstruction and subsequent allele-specific expression analysis^36,40,41^. We observed that the median number of SNVs per gene was two (**Supplemental Fig. 12a**) and the median distance between SNVs of the same gene was 511 nucleotides (**Supplementary Fig. 12b**). Using multi-SNV reads, haplotypes were elucidated by iteratively phasing SNVs along a given gene (**Methods**). Two haplotypes were reconstructed for 370 multi-SNV genes for which 94.6% of transcript reads had the majority of SNVs match a haplotype sequence. Thus, these haplotypes are generally representative of observed allele-specific transcripts.

Among the haplotypes, human leukocyte antigen (HLA) alleles were noteworthy given their diversity and function in adaptive immunity^42^. Consistent with their well-known polymorphism, a large number of SNVs were detected in the HLA genes, ranging from 46 in HLA-A to 8 in HLA-DRA (**Supplementary Fig. 12c**). We observed uneven mapping of transcript reads between the two alleles; HLA-DRA exhibited a striking 3.6-fold bias for transcripts mapping to haplotype 1 (H1) versus haplotype 2 (H2) (**Fig. 6a**). Imbalanced allele-specific expression is recognized as a pervasive feature of cancer, potentially stemming from alterations such as genomic structural variation and dysfunctional epigenetic regulation^43^. Here, in the context of HGSOC, we sought to systematically characterize the imbalanced allele-specific expression between tumor epithelial cells and the residual TME cell populations. We identified 33 genes displaying imbalanced allelic expression within the epithelial cell population but not in the remaining non-epithelial cells (Mann-Whitney *U* Test; Benjamini-Hochberg adjusted *P*<10^-6^ and *P*>0.05, respectively) (**Fig. 6b,c**). Among genes exhibiting epithelial-specific imbalanced allelic expression, VEGFA and CD276 are therapeutic targets for treatment of ovarian malignancies like HGSOC^44^ (**Fig 6b,d**). With scTaILoR-seq, phasing of SNVs permitted high-quality haplotype reconstruction and enabled quantitation of allele-specific expression among cellular populations within ovarian tumor samples.

**Fig. 6.**
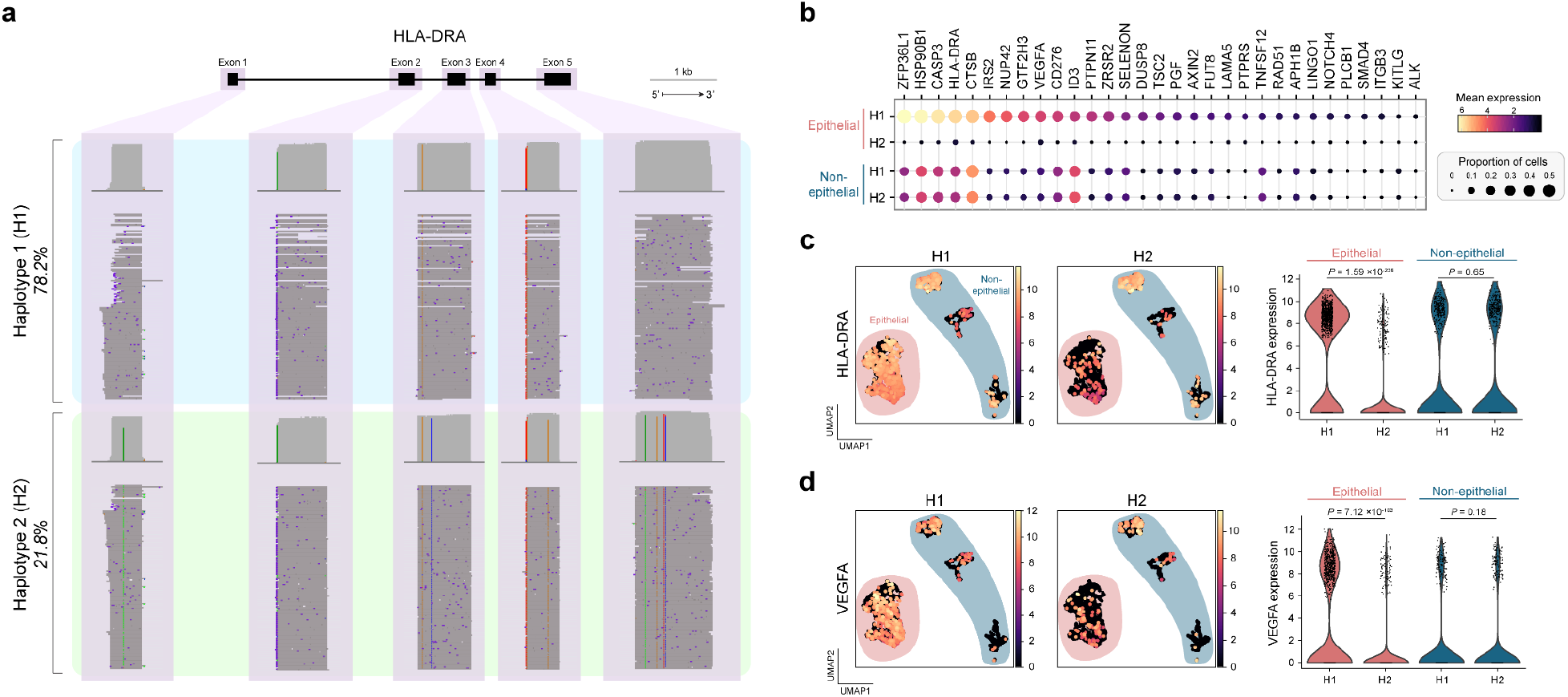
Variant phasing enables measurement of allelic imbalance at the single-cell level. **a**, Phased (H1 - blue, H2 - green) single-molecule read tracks for HLA-DRA. **b**, Allelic expression (H1/H2) and proportions of cells detected for epithelial and non-epithelial groups ranked by magnitude of imbalance. **c** and **d**, Plots for HLA-DRA and VEGFA, respectively: UMAP visualization of haplotype expression with epithelial cells highlighted in red and non-epithelial cells highlighted in blue. Violin plots show significant cell type-specific allelic imbalance (Mann-Whitney U test).

## Discussion

Recent improvements in nanopore sequencing chemistries, basecalling accuracy and bioinformatic tools have enabled single-cell long-read sequencing, which can deliver unprecedented insights into cell type-specific transcriptional diversity. However, there remain key challenges such as relatively lower throughput and template switch artifacts. Here, we developed scTaILoR-seq to partially address these challenges using an enrichment procedure that focuses on predetermined genes of interest while minimizing off-target artifacts. Because this method relies on designed gene panels, prior biological knowledge is required to optimally select hybridization probe sets. Fortunately, several expert-curated commercial probe panels–like the ones used in this study–are available for a range of biological applications. The use of custom probe panels further expands the scope of scTaILoR-seq to investigate specific biological systems and non-model organisms. This approach was developed to optimally allocate sequencing reads to hundreds or a few thousand genes of interest; whereas, current methods shallowly survey the whole transcriptome^18,45^ or deeply examine a narrow set of target genes^17,25^. With our approach, we demonstrate improved transcript detection sensitivity that enables a myriad of applications ranging from differential isoform expression analysis to discovery of sequence variants.

Fundamental to scRNA-seq is the ability to resolve reads by individual CBs and UMIs, which enables cell-specific quantitation of expression. Given the higher basecalling error rates of nanopore sequencing compared to short-read sequencing, prior long-read single-cell approaches have relied on supplemental short-read data to improve CB/UMI assignment accuracy^16,24^. Despite the inherent errors of nanopore reads, we demonstrate that scTaILoR-seq paired with the *wf-single-cell* workflow is capable of accurately producing single-cell transcriptomes from a complex tumor tissue without the need for supplemental short-read information.

Expansion of the scTaILoR-seq targeting panel to include immune-related genes permitted TCR repertoire profiling. Despite T cells comprising only 12% of the total cell population, scTaILoR-seq provided sufficient read depth at the TCR locus for highly specific and sensitive sequence reconstruction. While short-read TCR analysis typically requires specialized library preparation and/or 5’ RNA-seq, we demonstrate TCR reconstruction with scTaILoR-seq using conventional 3’ RNA-seq. Additionally, we were able to identify unique and expanded clonotypes which could provide insight into TME-specific T cell interactions and tumor antigens. Both the immune repertoires and the extent of clonal expansion are key determinants of the anti-tumor response and outcomes^46–48^.

Increased read depth and broader transcript coverage enabled more comprehensive detection of expressed SNVs, which was fundamental for the characterization of transcript structure alterations. Because many of these SNVs are proximal to annotated splice junctions, we suspect sequence-specific impacts on spliceosome function may contribute to the observed transcript structures. In some cases, a SNV and its associated site of structural divergence are within an A/T-rich region, which may be susceptible to internal priming during reverse transcription and/or second-strand synthesis^49^. While such artifacts would be considered false positives, we also observed opposing examples in which structural divergence was detected at non-A/T-rich regions and at A/T-rich sites that lacked structural divergence. Taken together, scTaILoR-seq enables the characterization of SNV-associated transcript structures which may be particularly impactful in evaluating the functional consequences of cancer mutations.

By reconstructing haplotypes from multi-SNV reads, we identified imbalanced allelic expression within tumor epithelial cells. Of potential therapeutic relevance is the observed allele-specific expression of VEGFA, which is the target of bevacizumab (Avastin) for treatment of platinum-resistant recurrent epithelial ovarian cancer^50,51^. Additionally, CD276 showed imbalanced allelic expression and is the target for clinical development as a cancer immunotherapy^44^. Thus, beyond the fundamental biological insights enabled by scTaILoR-seq, the ability to simultaneously characterize cell- and allele-specific transcriptional variation has the potential to impact diagnostic and therapeutic approaches.

While we have highlighted many capabilities of scTaILoR-seq, there are several features that can be developed to further improve this method. First, the analyses presented here focused primarily on annotated transcripts. It remains to be seen what effect gene targeting has on novel isoform discovery. Second, abortive reverse transcription hinders long-read analysis due to truncated reads and may be addressed with alternative reverse transcription strategies. Finally, we expect that this approach can be adapted to many of the emerging and existing commercial scRNA-seq platforms (*e.g.* droplet, nanowell, and combinatorial indexing) in addition to synergistic technologies like spatial transcriptomics. With these adaptations in mind, scTaILoR-seq provides an attractive option for efficient exploration of full-length transcriptomes, especially for large-scale single-cell atlasing initiatives.

We developed scTaILoR-seq, a novel targeted long-read sequencing approach for single-cell characterization of transcriptional variation. scTaILoR-seq efficiently allocates sequencing throughput to improve detection and quantitation of transcripts of interest at multiple resolutions: from exon structure down to single nucleotide variants.

## Methods

### Single-cell isolation and 10X Genomics 3’ cDNA generation

#### Cell Lines

To evaluate sensitivity and the robustness of our method we obtained three ovarian cancer cell lines SK-OV-3, IGROV1 and COV504. SK-OV-3 and IGROV1 were maintained in RPMI-1640 medium, supplemented with 10% fetal bovine serum (FBS) and 2 mM L-Glutamine. COV504 cells were maintained in DMEM supplemented with 10% FBS and 2 mM L-Glutamine. Cells from each cell line were prepared following the 10x Genomics Cell Preparation Guide (CG000053_CellPrepGuide_RevC) and combined at equal cell concentrations prior to loading onto the 10x Genomics Chromium Controller at a concentration of 1000 cells/μL. cDNA generated through the single-cell platform was then split for single-cell targeted long-read enrichment (see below) or for generating scRNA-seq short-read libraries using the 10x v3.1 protocol (CG000204_ChromiumNextGEMSingleCell3’v3.1_RevD).

#### Dissociated Tumor Cells

Ovarian cancer dissociated tumor cells were purchased through Discovery Life Sciences (Huntsville, AL). Both samples were stage III, treatment naive with cancer subtypes Clear Cell Carcinoma and High Grade Serous Carcinoma. All clinical metadata was de-identified and samples were IRB compliant. Cells were thawed and prepared following the recommended 10x Genomics Cell Preparation Guide shown above with minor adjustments. Cells were thawed for 2 minutes and placed into 15 mL of warm RPMI media containing 10% FBS media. Cells were spun at 300g for 5 minutes. DNase I was added after the first spin to prevent clumping. Three additional spins were performed with 1X PBS with 0.04% BSA to ensure proper removal of DNase I prior to 10x loading. Cells were counted and checked for viability using Vi-Cell XR (Beckman Coulter). The viability was 88.3% and 82.5% and the target capture was for 6000 cells prior to injection. Both the cell lines and primary tumor cells were run on Chip G using the 10x v3.1 kit for generating the cDNA (CG000204_ChromiumNextGEMSingleCell3’v3.1_RevD). The cDNA amplification step was modified by extending the elongation time to 2 minutes rather than the recommended 1 minute. cDNA generated through the droplet single-cell platform was then split for either long-read enrichment or for preparing scRNA-seq short-read libraries using the 10x v3.1 protocol (CG000204_ChromiumNextGEMSingleCell3’v3.1_RevD).

### Illumina library generation and sequencing

Whole transcriptome short-read libraries were dual-indexed and sequenced paired-end on the Illumina NovaSeq 6000 p with the recommended 10x run parameters (Read 1 - 28 cycles, i7 - 10 cycles, i5 - 10 cycles and Read 2 - 90 cycles). Targeted short-read libraries were dual-indexed and sequenced paired-end on the Illumina NextSeq 2000 following the same run parameters as shown above.

### Single-cell targeted gene enrichment for long-read sequencing

#### Pre-Amplification

To be able to get the recommended 300 ng of input for gene enrichment, approximately 10 ng of the 10x cDNA derived from the cell lines were split into two reactions and amplified an additional 5 cycles of PCR using two customized primers: (1) TruSeq Read 1 forward primer 5’ (Fwd_partial_read1) and (2) partial TSO reverse primer (Rev_partial_TSO) (**Supplementary Table 3**). The PCR reaction was carried out using 2X LongAmp Taq (NEB) with the following PCR parameters 94°C for 3 minutes, with 5 cycles of 94°C 30 seconds, 60°C 15 seconds, and 65°C for 3 minutes, with a final extension of 65°C for 5 minutes. The cDNA was then purified using 0.8X SPRI beads to remove unwanted primers and eluted in 30 μL H_2_O.

#### R2C2

Post-enriched cDNA was used for input into an R2C2 reaction following the protocol previously described^19^. Briefly, 100 ng of the targeted cDNA was circularized using Gibson assembly (NEBBuilder HiFi DNA assembly mix) with a custom splint that is compatible with 10x cDNA containing both the Read1 (10X_UMI_Splint_Forward) and TSO sequences (10X_UMI_Splint_Reverse) (Supplementary Table 3). Any non-circularized byproducts were then digested using an exonuclease mixture of Lambda, Exo I and Exo III (NEB) and incubated at 37°C overnight. Post overnight digestion the reaction was cleaned up with 0.8X SPRI and eluted in 30 μL. The circularized product was separated into three different reactions and amplified using rolling circular amplification using Phi29 (NEB) and incubated at 30°C overnight. To debranch the Phi29 product, a T7 endonuclease (NEB) digestion was performed and incubated at 37°C on a thermal shaker at 1000 RPM for 2 hours. A final 0.5X SPRI purification is performed to enrich longer molecules >500 bp, about 1 μg should be recovered.

#### Cell Lines

The pan-cancer gene panel (*n*=1,253 genes) was designed by 10x Genomics containing 120 bp probes tiled across known annotated exons covering both sense and antisense strands. The cDNA hybridization using the pre-designed panels was performed following the 10x protocol (CG000293_TargetedGeneExpression_SingleCell_UG_RevF) with minor changes. We incorporated TSO blockers (1 μM) during Step 1.1 in the pre-hybridization pooling and drying step (Supplementary Table 3). The pre-hybridization was carried out using 300 ng of cDNA, 20 μL of COT DNA, 0.8 μL of TSO blockers, and 2 μL of Universal Blockers. The samples were dried using the SpeedVac Savant DNA120 concentrator (Thermo Fisher Scientific) on ‘Medium’ setting. Following the hybridization, 5 cycles of PCR were performed using the same cDNA primers described in the Pre-Amplification step (1) Fwd_partial_read1 and (2) Rev_partial_TSO to amplify molecules off the bead. The following PCR conditions were the same as described in the Pre-Amplification step.

#### Dissociated Tumor Cells

For the primary tumor samples the gene enrichment was performed as discussed above with the exception that the same cDNA was separated into two enrichments one using the pan-cancer gene panel (*n*=1,253 genes) and the other with the Immune gene panel (*n*=1,056 genes). The samples were targeted following the 10x protocol (CG000293_TargetedGeneExpression_SingleCell_UG_RevF) with minor changes as indicated above incorporating TSO blockers. Following the hybridization, 5 cycles of PCR were performed using the non-biotinylated primers (1) Fwd_3580_partial_read1_defined and (2) Rev_PR2_partial_TSO_defined from the single-cell ONT protocol (Supplementary Table 3, single-cell-transcriptomics-10x-SST_v9148_v111_revB). The PCR reaction was carried out using 2X LongAmp Taq (NEB) with the following PCR parameters 94°C for 3 minutes, with 5 cycles of 94°C 30 seconds, 60°C 15 seconds, and 65°C for 3 minutes, with a final extension of 65°C for 5 minutes. The post cDNA hybridized product was then purified with 0.8X SPRI beads to remove unwanted primers and eluted in 40 μL of H_2_O. cDNA concentration was measured using Qubit dsDNA HS kit and the size distribution analyzed using Tapestation D5000 Screen Tape (Agilent Technologies).

#### TSO Artifact Mitigation

Post hybridization artifact mitigation was performed using the biotinylated version of the forward primer from the ONT protocol, [Btn]Fwd_3580_partial_read1_defined (**Supplementary Table 3**). The PCR reaction was carried out using 2X LongAmp Taq (NEB) with the following PCR parameters 94°C for 3 minutes, with 3 cycles of 94°C 30 seconds, 60°C 15 seconds, and 65°C for 3 minutes, with a final extension of 65°C for 5 minutes. Full-length cDNA was captured using 15 μL M270 streptavidin beads (Thermo Fisher Scientific) that were washed three times with SSPE buffer (150 mM NaCl, 10 mM NaH_2_PO_4_, and 1 mM EDTA) and resuspended in 10 μL of 5X SSPE buffer (750 mM NaCl, 50 mM NaH_2_PO_4_, and 5 mM EDTA). The cDNA obtained from the gene enrichment step was combined with 10 μL M270 beads and incubated at room temperature for 15 minutes. After incubation, the cDNA-bead conjugate was washed twice with 1 mL of 1X SSPE. A final wash was performed with 200 μL of 10 mM Tris-HCl (pH 8.0) and the beads bound to the sample were resuspended in 10 μL H_2_O. A final PCR was performed on-bead using the cDNA primers (cPRM) from the SQK-PCS111 kit following the PCR conditions from the single-cell ONT protocol (single-cell-transcriptomics-10x-SST_v9148_v111_revB). The cDNA was cleaned up with 0.8X SPRI and eluted in 15 μL. The concentration and quality of the sample was evaluated with Qubit dsDNA HS kit and Tapestation D5000 Screen Tape (Agilent Technologies). The expected recovery was above 50 ng.

### ONT library preparation and nanopore sequencing

#### Cell Lines

For the mixed ovarian cell lines, library preparation for nanopore sequencing was performed according to the LSK-109 protocol (ONT). For the targeted mixed ovarian cell line samples, the final libraries (targeted, targeted+AM and targeted+R2C2) were loaded onto a total of seven MinION flowcells (FLO-MIN106D). Approximately 25-30 fmol of the library was loaded for each run. The samples were sequenced for 72 hours and basecalled using Guppy v6.0.1. For the un-targeted sample, library preparation was performed according to the LSK-110 protocol. A total of 125 fmol was loaded onto a single PromethION flowcell (FLO-PR002), sequenced for 72 hours and basecalled using Guppy v6.0.1.

#### Dissociated Tumor Cells

After post enrichment and artifact mitigation the rapid adapter addition was performed following SQK-PCS111 protocol. Final libraries (125 fmol per library) across both patient samples were loaded onto a total of 4 PromethION flowcells (FLO-PRO002). The samples were sequenced for 72 hours and basecalled using Guppy v6.0.1.

### Long-read CB and UMI assignment

‘SiCeLoR’ (https://github.com/ucagenomix/sicelore/commit/b057aa0f7948d2e8f64140b8ec99c2f3bb4b6d53) was used with default settings to process reads when companion short-read data were available. When considering complete reads, 79.5% (average of 2 replicates) could be assigned to a known cell barcode Of those, approximately 68% were matched to UMIs identified from short-read sequencing. These values are consistent with recent single-cell nanopore long-read sequencing efforts. Next, for CB/UMI assignment without companion short-read data, we used ‘wf-single-cell’ (https://github.com/epi2me-labs/wf-single-cell; v0.1.5) with default settings. UMI-deduplication of the resultant tagged bam file was performed using ‘UMI-tools’ (v1.1.0) with the following settings for both ‘group’ and ‘dedup’ functions: --per-cell --per-gene --extract-umi-method=tag --umi-tag=UB --cell-tag=CB --gene-tag=GN. In general, we used the GRCh38 human reference genome and GENCODE v32/Ensembl 98 annotations provided by 10x Genomics (2020-A; July 7, 2020; https://support.10xgenomics.com/single-cell-gene-expression/software/downloads/latest).

### Single-cell data analysis

Cell-by-transcript count matrices were generated directly from ‘SiCeLoR’ or from ‘IsoQuant’ (v3.1.0) with default settings using CB-tagged bam files generated by ‘wf-single-cell‘. The count matrices were processed using ‘scanpy’ (v1.9.1) as follows: 1) normalize counts per cell (target_sum=10^6^), 2) log1p transform, and 3) scale to unit variance and zero mean. Unsupervised clustering of cell subgroups was performed using the Leiden algorithm applied to the neighborhood graph of principal components. Differential expression of both genes and transcripts computed using Welch’s t-test (method=”t-test_overestim_var”). Geneset expression scores were calculated using the ‘score_genes’ function from ‘scanpy‘. Pathway activity scores were calculated using the ‘progeny’ function (z_scores=TRUE, organism=“Human”, top=300, perm=100) from ‘PROGENy’ (v1.18.0). For genetic-deconvolution of cell identity, we used ‘souporcell’ (v2.0) with known genotypes provided as a ‘BCFtools‘-merged ‘Clair3‘-derived .vcf file. Cell multiplets were identified using ‘Scrublet’ (v0.2.3), implemented within ‘scanpy’ with default settings. To integrate the expression matrices from the pan-cancer and immune panels, we applied a scalar offset. From the transcripts of 258 genes shared between the two panels, the scalar offset was computed as the mean slope of 10-fold cross-validated (CB-shuffled) linear regression slopes (‘sklearn’ v1.0.1) using mean transcript expression (cell count-normalized and log1p-transformed). Subsequently, single-cell transcript expression values corresponding to the response variables were multiplied by the scalar offset. To construct the integrated expression matrix, scaled transcript expression values private to the response variables were joined (CB-matched) to the expression matrix corresponding to the predictor variables.

### T cell receptor reconstruction

The immune enrichment panel design comprises probes targeting the constant TCR genes: TRAC, TRDC, TRBC2, and TRGC1. The TRBC1 and TRGC2 genes were not included in the panel as they have high homology to selected probes. scTaILoR-seq reads were processed by *TRUST4* using the parameters *“--ref human_IMGT+C.fa --barcode CB --UMI UB.”*

### Variant analysis

Cell subpopulation reads were aggregated from CB-tagged bam records using ‘pysam’ (v0.16.0.1), then variants were called using ‘Clair3’ (v0.1-r11) with pretrained model r941_prom_hac_g360+g422 (--platform=ont --enable_phase --fast_mode). For analysis of variants associated with transcript structural divergence, Clair3-derived variant calls were filtered (DP>=100 and QUAL>=15). Variant calls were scored for cryptic splicing using ‘SpliceAÌ (v1.3.1, -D 500). Then, for each variant, aligned reads were partitioned by observed base matching REF or ALT values (*via* ‘pysam‘). Read coverage of resultant REF- and ALT-specific bam files were computed using ‘bamCoverag’ (v3.5.0, --binSize 1). The Pearson correlation coefficient (*r*) between REF- and ALT-specific read coverage was calculated (minimum depth>=50). The degree of transcript structural divergence was defined as the variance unexplained (1-*r*^2^). For variants exhibiting non-zero coverage divergence, linear regression residuals between REF- and ALT-specific coverage at single-base resolution were mapped to annotated transcript structural features: CDS and UTR/Intron. Then, the proportion of bases with residual z-score>0.5 within each structural feature was max-normalized per variant before agglomerative hierarchical clustering (method=“ward”, metric=“euclidean”) using ‘SciPy’ (v1.7.3).

### Haplotype analysis

Only reads with at least two detected SNVs were considered for haplotype reconstruction. For each gene, the observed variant status of each read was encoded as a vector of position-sorted SNV sites (*n*=number of detected SNVs within the gene) with the following values: undetermined=0, REF=1, ALT=2. The SNV vector with the highest read count was used as the seed haplotype. For each element in this seed vector equal to 0 (*i.e.* undetermined), the variant status was determined as follows:

1. Identify all reads that contain at least one determined SNV site (REF=1 or ALT=2) from the current SNV vector in addition to the undetermined site.
2. Update variant status at undetermined site based on highest frequency nucleotide identity (REF or ALT) at that position.

Haplotype reconstruction was complete when all SNV sites were determined. Then, allele-specific reads with a majority of SNVs (>50%) matching the haplotype were masked before a second haplotype was determined as outlined above. The final allele-specific read annotations were similar to above (*i.e.* majority of haplotype-matching SNVs per read) but omits SNV sites with shared identity between H1 and H2.

### Data and code availability

Sequencing data were deposited to NCBI Sequence Read Archive (SRA) under the BioProject accession PRJNA993664. Code to be made available upon reasonable request.

## Acknowledgements

We would like to thank Chris Rose, Carolina Galan, Andrew McKay, and Spyridon Darmanis for critical review of the manuscript. We would also like to thank Oleg Mayba and Bing Wu for helpful discussions regarding analysis.

## Contributions

A.B. and W.S. conceived the study. K.S. provided ovarian cell lines for single-cell RNA-seq. H.M. assisted with single-cell RNA-seq. A.B., W.S. and J.L. performed long-read sequencing. N.P., J.L., A.X-M., M.S. performed short-read sequencing. D.L., A.B., W.S. and T.S-W. performed analysis. Z.M. and W.S. supervised the project. W.S., D.L., Z.M. and A.B. wrote the manuscript. All authors discussed the results and approved the manuscript.

## Corresponding authors

Correspondence to Zora Modrusan and William Stephenson.

## Ethics declarations

All authors are employees and shareholders of Genentech.

## Supplementary Note

### Short-read untargeted and targeted sequencing

We performed droplet-based single-cell 3’-end RNA sequencing on a mixture of three human ovarian cancer lines (SK-OV-3, COV504, and IGROV-1) and generated a whole transcriptome short-read library (while reserving full-length cDNA) in addition to performing targeted gene enrichment using the 10x Genomics Pan-Cancer panel comprising 1,253 genes. We performed short-read sequencing on the untargeted and targeted libraries to depths of 796M and 173M reads representing 62.2% and 89.5% sequencing saturation, respectively. Target enrichment recovered 85.2% of barcodes found within the untargeted approach (“intersecting cell barcodes” - orange); notably we observed that missing cell barcodes (“non-intersecting cell barcodes” - blue) represented lower quality cells in the untargeted dataset with lower counts and higher mitochondrial content (see figure below). These cells were likely excluded due to distinct cell barcode/UMI cutoffs used in the targeted approach (https://support.10xgenomics.com/single-cell-gene-expression/software/pipelines/latest/algorithms/targeted#targeted-cell-calling). In addition to running Cell Ranger `count` on untargeted and targeted libraries individually (targeted performed including the `--target-panel` path option), further analysis was performed using Cell Ranger `targeted-compare`. Sequencing the untargeted and targeted libraries resulted in 4% and 82.1% of reads confidently mapped to the targeted transcriptome respectively, representing a 17.7-fold mean read enrichment.

**Supplementary Fig. 1.**
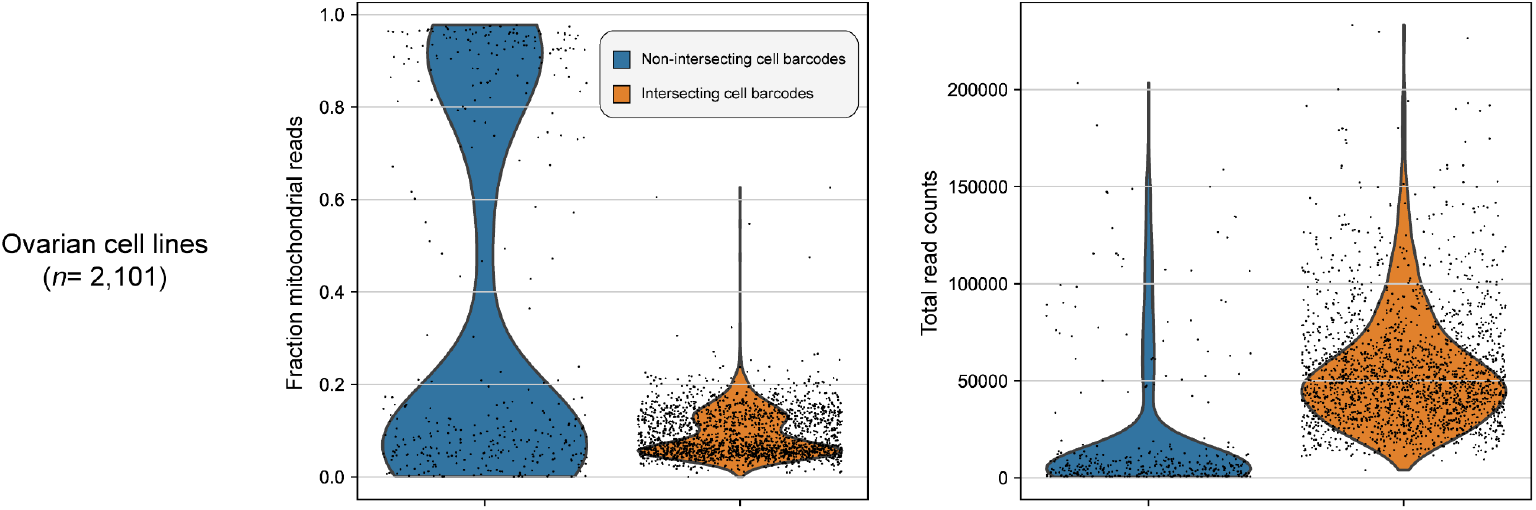
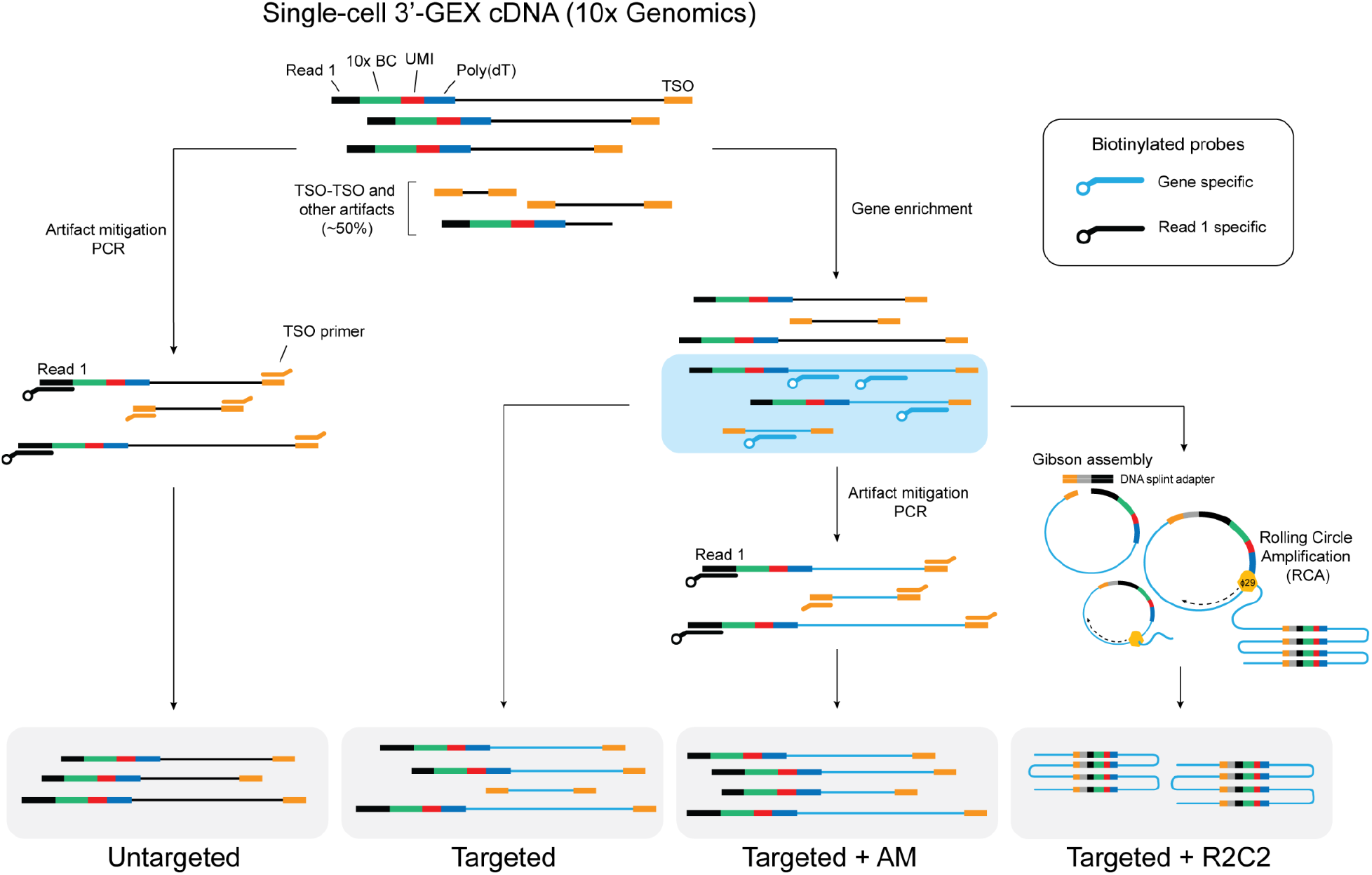
Long-read enrichment approaches. Schematic detailing enrichment optimization approaches for long read sequencing. The untargeted approach contains an artifact mitigation step (+AM) using biotinylated probes against R1 (read 1) primer. The targeted approach uses biotinylated probes for enriching genes of interest only. The targeted+AM performs both gene enrichment and artifact mitigation and the targeted+R2C2 approach utilizes gene enrichment combined with a rolling-circle amplification based library preparation method for read consensus.

**Supplementary Fig. 2.**
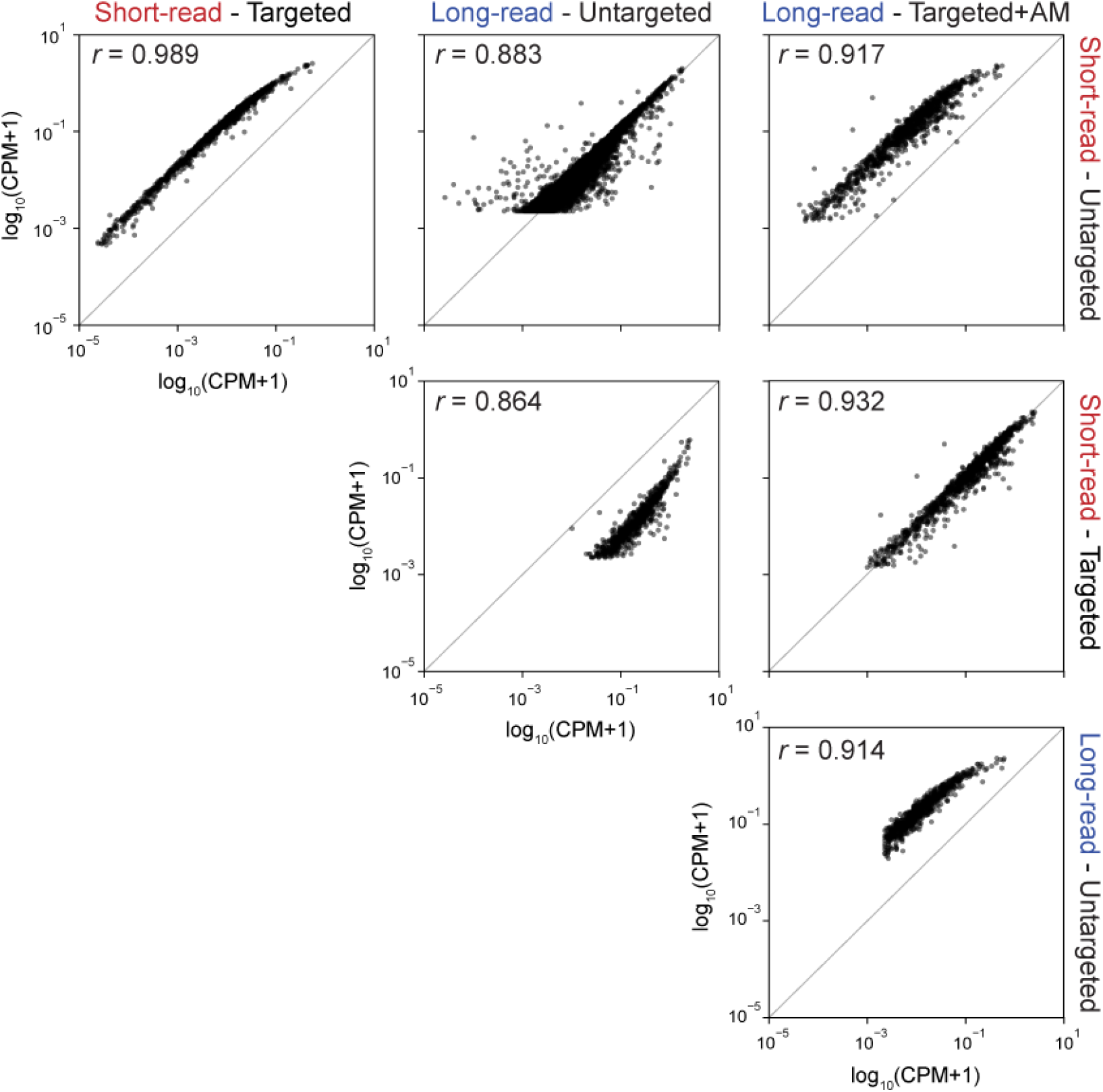
Mean gene expression correlations between methods. Scatter plot grid containing pairwise comparisons at the pseudo bulk level across three enrichment approaches in combination with short and long read sequencing. The x-axes are indicated by the row labels at right and the y-axes are indicated by the column labels at top. Correlations of mean gene expression are given as log10 (CPM+1). Pearson *r* is given for each gene across methods.

**Supplementary Fig. 3.**
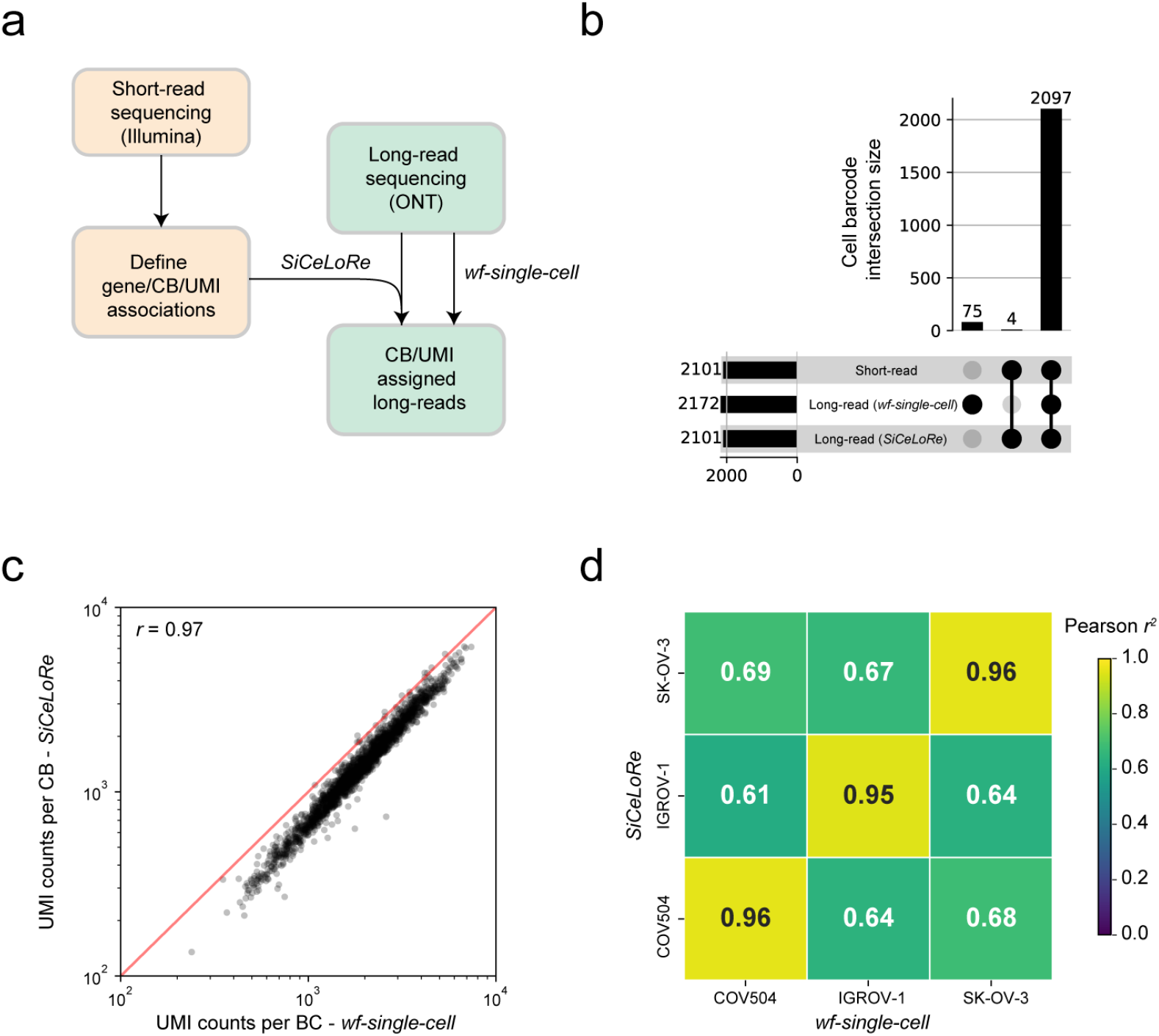
Analysis of guided and unguided CB/UMI demultiplexing approaches. **a**, Schematic of guided (*i.e. SiCeLoRe*; orange) and unguided (*i.e. wf-single-cell*; green) approaches. **b**, Upset plot showing overlap of detected CBs across guided, unguided and short-read approaches. **c**, Scatter plot showing UMI counts per CB for unguided vs guided approaches. **d**, Heatmap showing Pearson correlation results for unguided vs guided approaches with respect to cell line-specific mean gene expression. Tile color represents the squared Pearson correlation coefficient.

**Supplementary Fig. 4.**
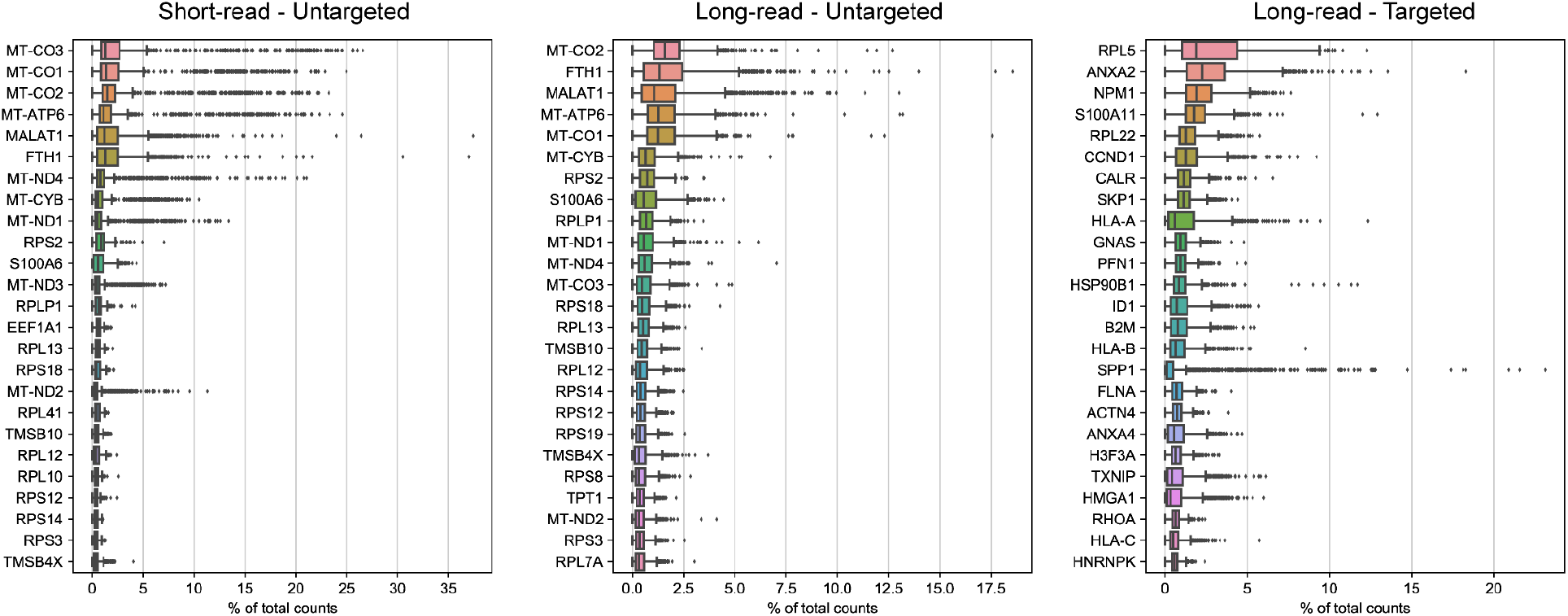
Comparison of top-25 highly expressed genes. Short-read untargeted, long-read untargeted, and long-read targeted boxplots showing top-25 highly expressed genes (y axis) calculated as percent total counts (x-axis).

**Supplementary Fig. 5.**
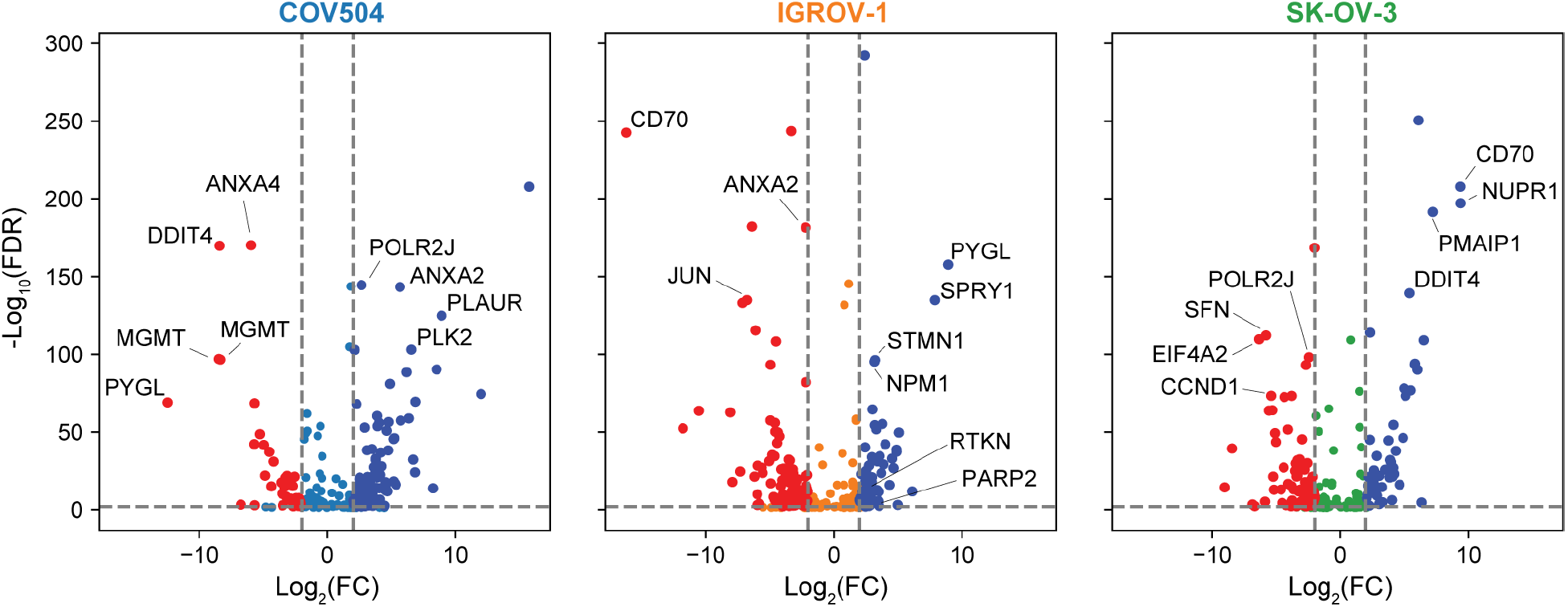
Differential transcript expression among three ovarian cancer cell lines. Volcano plots showing differential transcript expression for each cell line vs. rest: **left**, COV504 vs. IGROV-1 and SK-OV-3. **center**, IGROV-1 vs. COV504 and SK-OV-3. **right**, SK-OV-3 vs. COV504 and IGROV-1. Only gene names have been used to annotate individual transcripts in the plots. Significantly up-regulated transcripts are colored dark blue and down-regulated transcripts are colored red. The x-axis units are log base 2 fold-change and the y-axis units are negative log base 10 false discovery rate (Benjamini-Hochberg multiple testing adjustment). Dashed lines represent 2 fold-change and *P*<0.05 significance thresholds.

**Supplementary Fig. 6.**
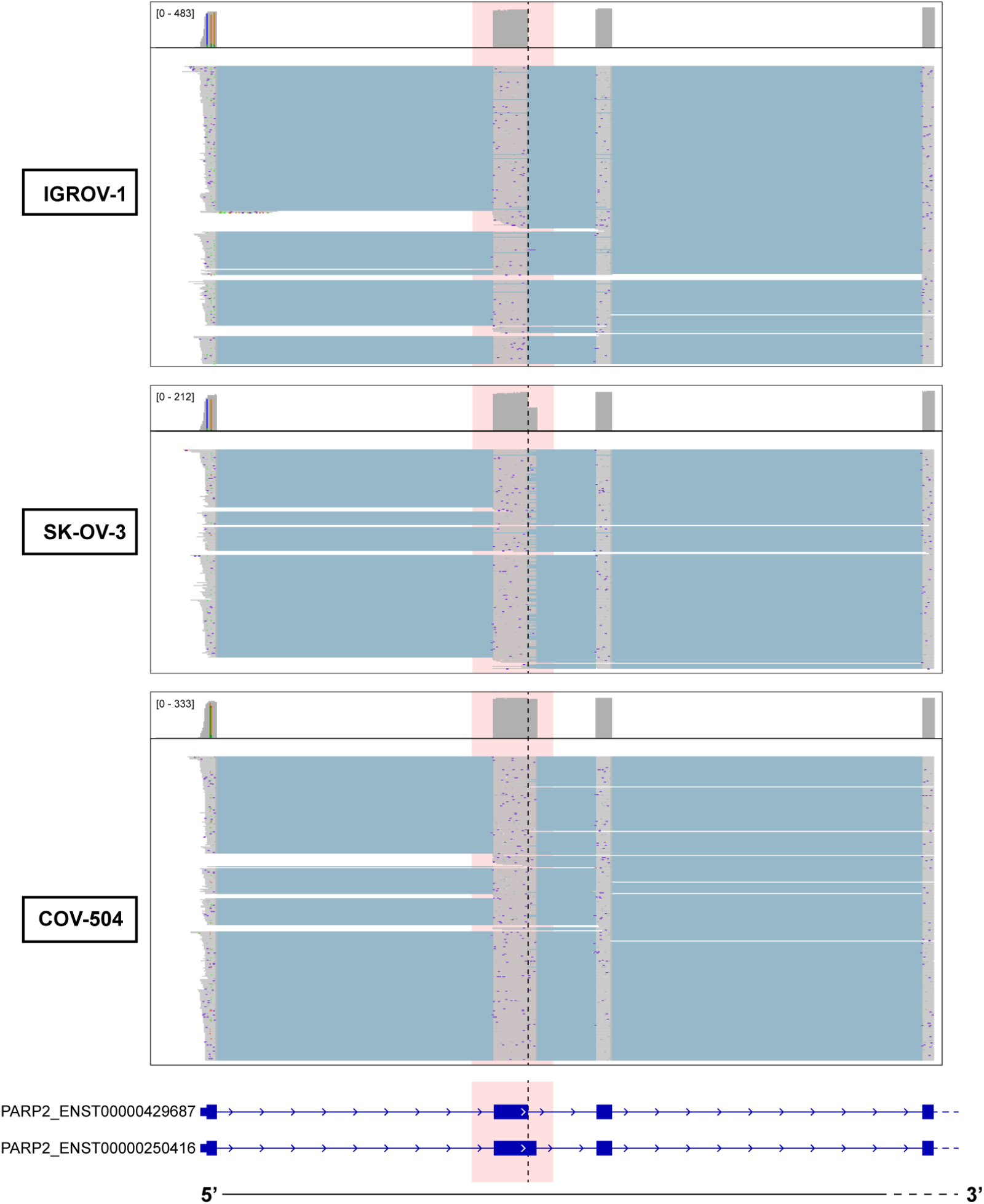
PARP2 differential transcript usage among cell lines. IGV tracks showing reads, segregated by cell line, aligned to PARP2. Two transcript models are shown below the alignment tracks. The pink rectangle highlights structural variation of exon 2 and its differential usage among the three cell lines.

**Supplementary Fig. 7.**
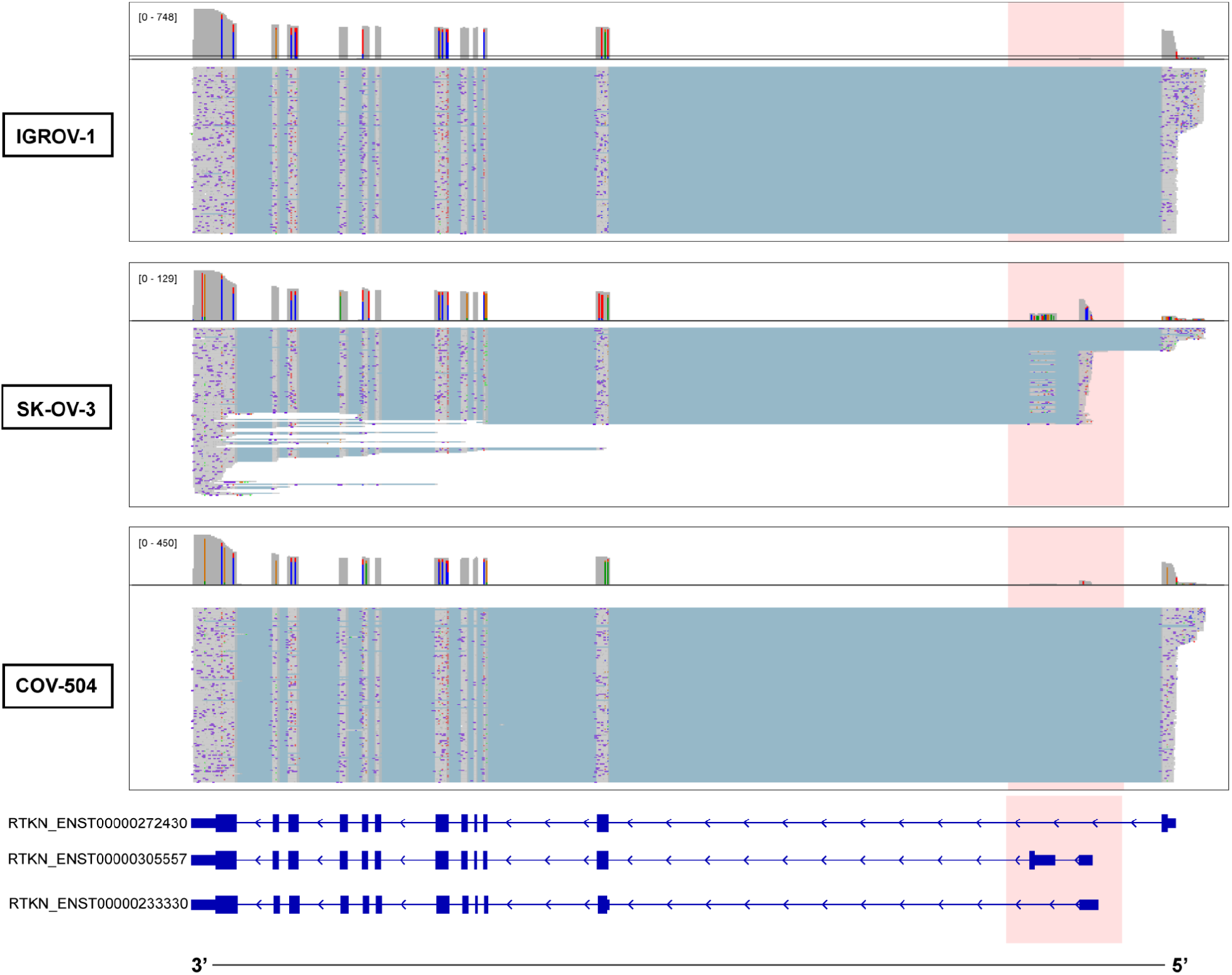
RTKN differential transcript usage among cell lines. IGV tracks showing reads, segregated by cell line, aligned to RTKN. Three transcript models are shown below the alignment tracks. The pink rectangle highlights structural variation of the 5’-UTR and first exon and its differential usage among the three cell lines.

**Supplementary Fig. 8.**
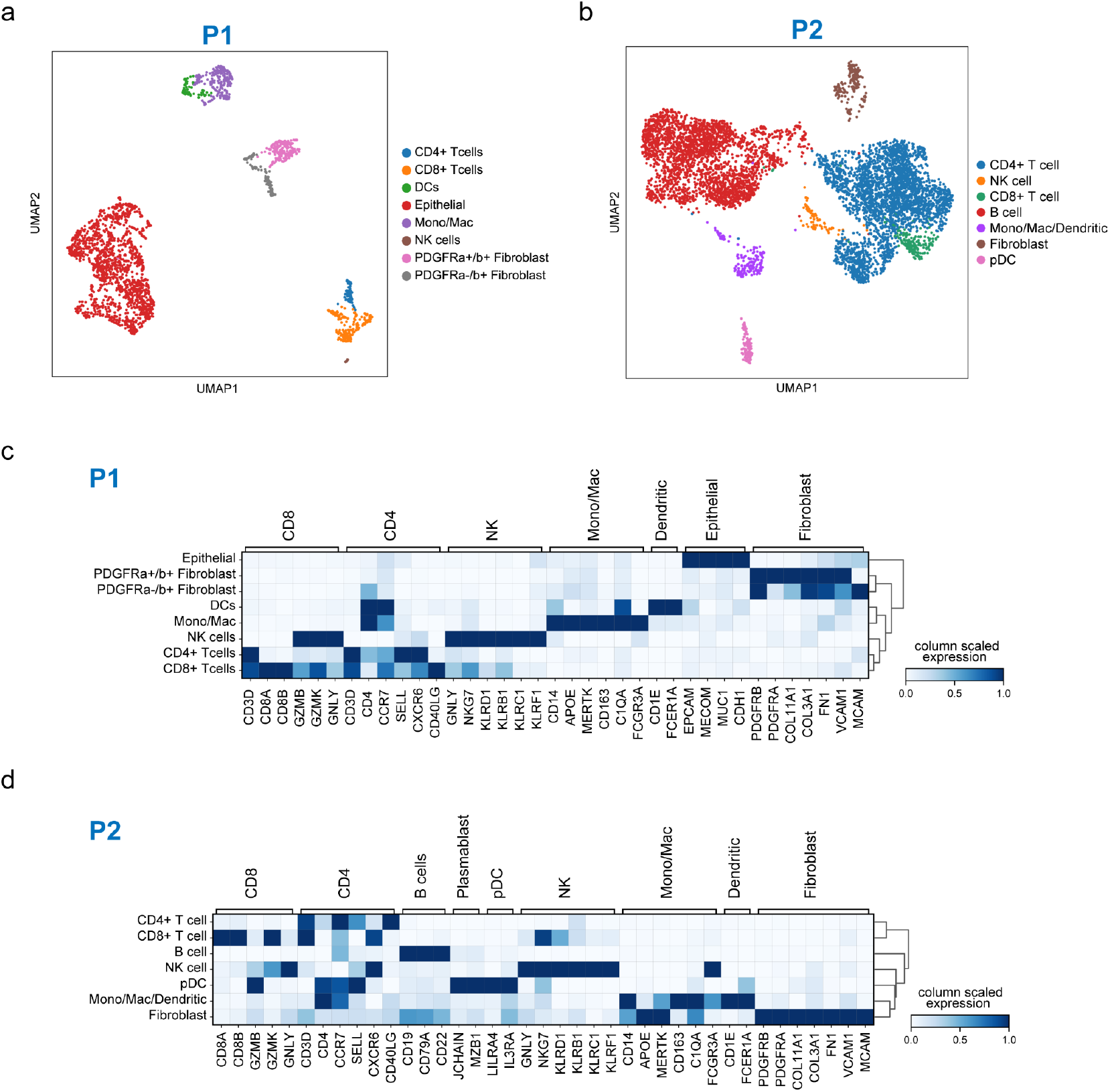
Patient 1 (P1) and Patient 2 (P2) cell annotations and expression of canonical marker genes. a, Gene-level UMAP projection of cells profiled from P1 processed with scTaILoR-seq workflow, colors represent annotated cell types. **b,** Gene-level UMAP projection of cells profiled from P2 processed with scTaILoR-seq workflow, colors represent annotated cell types. **c**, Matrix plot showing mean expression of canonical marker genes grouped by cell types found within P1. The tile color represents a scaled expression from lowest (0) to highest (1) mean expression. **d**, The same representation as in c for P2.

**Supplementary Fig. 9.**
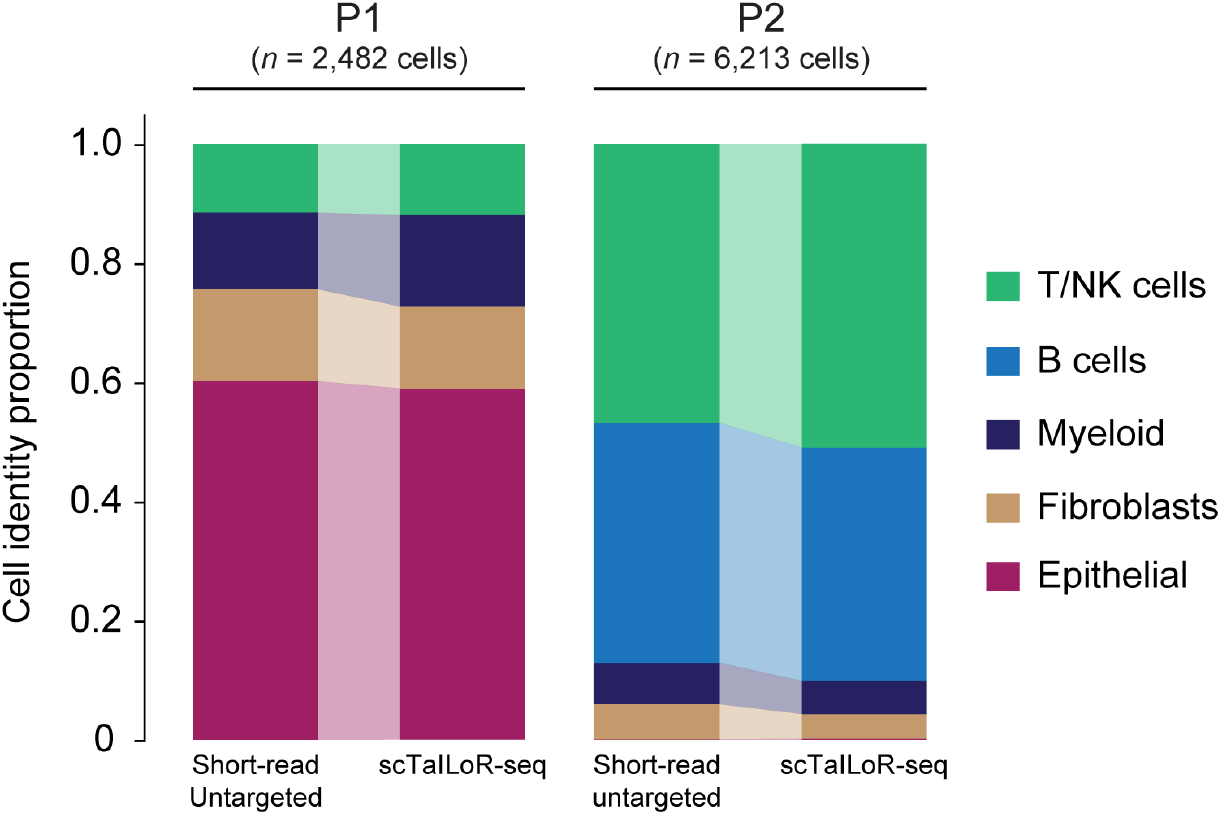
Major cell type proportions discerned with short- and long-read (scTaILoR-seq) sequencing across patient samples. Barplot showing cell type proportions across major cell types found within patient samples, P1 (High-grade serous ovarian cancer) and P2 (Clear Cell Carcinoma).

**Supplementary Fig. 10.**
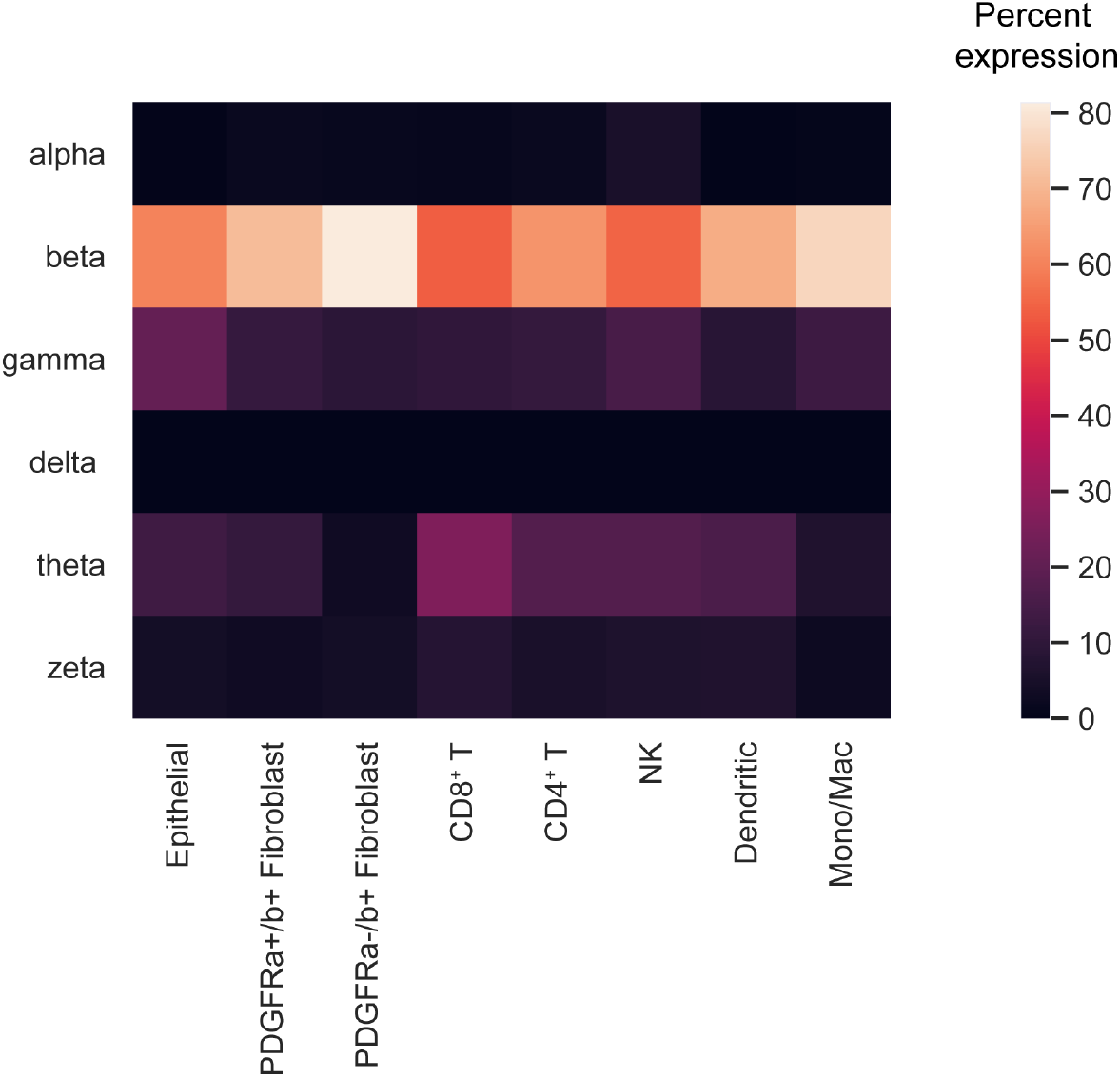
Cell type-specific IL32 differential transcript usage. Heatmap showing relative mean IL32 isoform expression (y-axis) scaled per cell type (x-axis). The tile color represents mean isoform expression relative to total IL32 expression within each cell type (*i.e.* percent expression; see legend).

**Supplementary Fig. 11.**
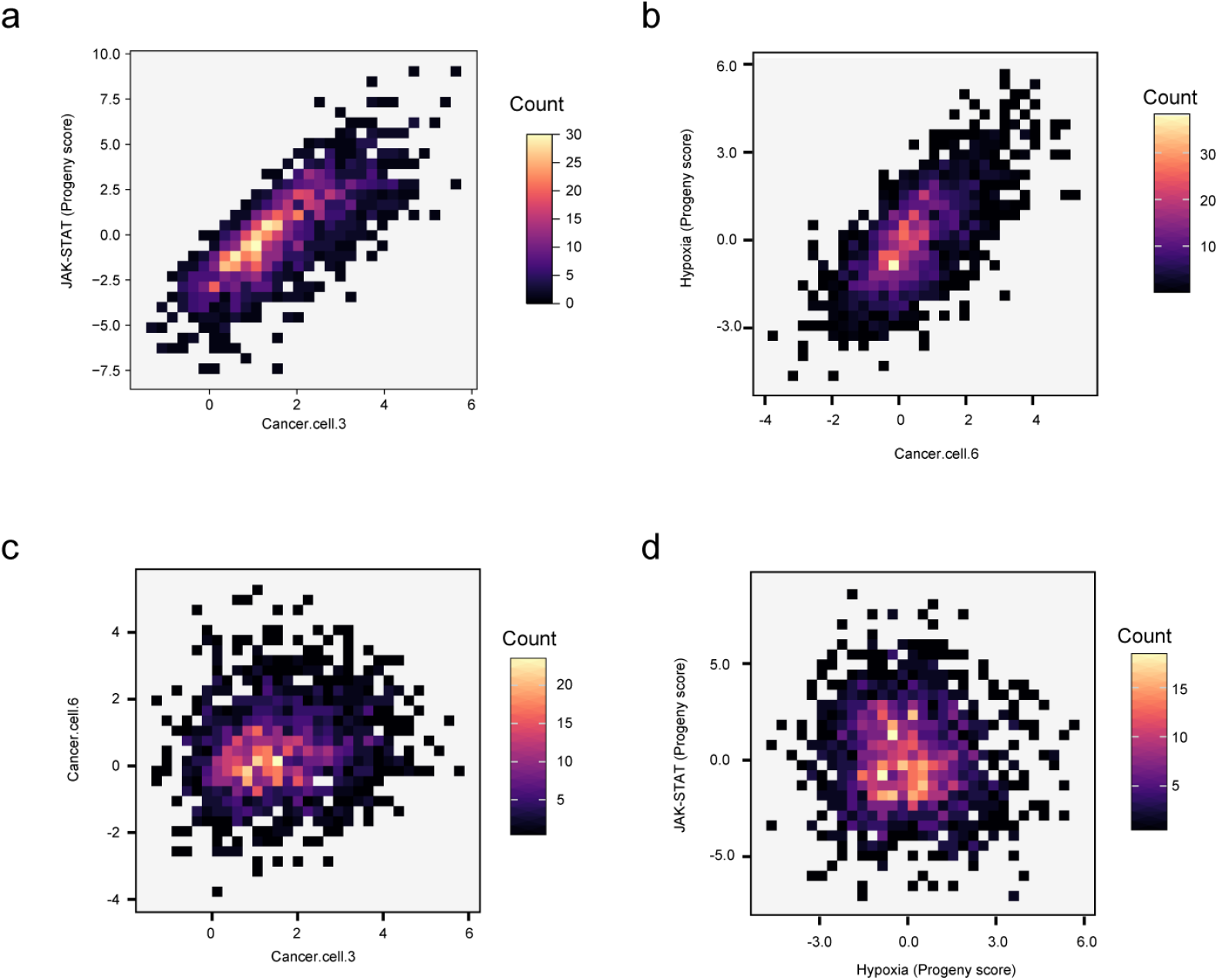
Single-cell correlations of PROGENy pathway activities and cancer-associated gene signatures. 2D bin plots with each tile color-coded by cell count for the following correlations: **a**, MSK Cancer.cell.3 expression signature vs. PROGENy JAK-STAT pathway activity (Pearson *r*=0.79). **b**, MSK Cancer.cell.6 expression signature vs. PROGENy Hypoxia pathway activity (Pearson *r*=0.68). **c**, MSK Cancer.cell.3 expression signature vs. MSK Cancer.cell.6 expression signature (Pearson *r*=0.15). **d**, PROGENy Hypoxia pathway activity vs. PROGENy JAK-STAT pathway activity (Pearson *r*=-0.08).

**Supplementary Fig. 12.**
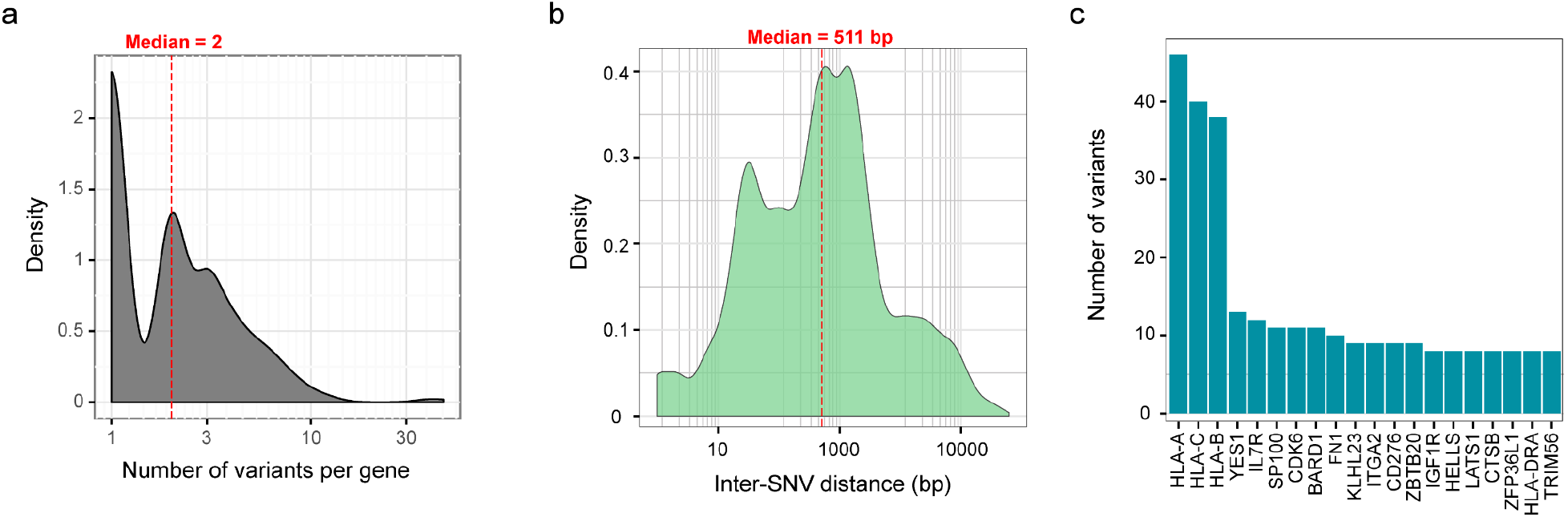
Characterization of SNV frequency and distribution. **a**, Density distribution of detected SNVs per gene. **b**, Density distribution of distance between detected SNVs. **c**, Top-20 genes based on number of detected SNVs.

**Supplementary Table 1.** Sequencing approaches and QC metrics. Read metrics across short- and long-read sequencing experiments. Enrichment strategy used, flow cell types, library preparation kit, passing reads and N50 are shown.

| Sample | Seq Type | Enrichment | Strategy | Flow Cell | Library Prep | Passing Reads (M) | N50 |
| --- | --- | --- | --- | --- | --- | --- | --- |
| Mixed Ovarian | ONT | No | Un-targeted | FLO-PRO002 | LSK-110 | 89.81 | 1420 |
| Mixed Ovarian | ONT | Yes | Pan Cancer Targeted | FLO-MIN106D | LSK-109 | 9.54 | 1440 |
| Mixed Ovarian | ONT | Yes | Pan Cancer Targeted | FLO-MIN112 | SQK-Q20EA | 9.60 | 1540 |
| Mixed Ovarian | ONT | Yes | Pan Cancer Targeted +AM | FLO-MIN106D | LSK-109 | 14.56 | 1230 |
| Mixed Ovarian | ONT | Yes | Pan Cancer Targeted +AM | FLO-MIN106D | LSK-109 | 9.54 | 1440 |
| Mixed Ovarian | ONT | Yes | Pan Cancer Targeted + R2C2 | FLO-MIN106D | LSK-109 | 4.53 | 4410 |
| Mixed Ovarian | ONT | Yes | Pan Cancer Targeted+ R2C2 | FLO-MIN106D | LSK-109 | 5.32 | 4680 |
| Mixed Ovarian | ONT | Yes | Pan Cancer Targeted+ R2C2 | FLO-MIN106D | LSK-109 | 3.55 | 5010 |
| Mixed Ovarian | ONT | Yes | Pan Cancer Targeted+R2C2 | FLO-MIN106D | LSK-109 | 3.25 | 5200 |
| Mixed Ovarian | Illumina | Yes | N/A | P1 300 cycle | 10X Lib Prep | 60.65 | N/A |
| Mixed Ovarian | Illumina | No | N/A | S2 | 10X Lib Prep | 314.5 | N/A |
| DTC_Patient 1 | ONT | No | N/A | FLO-PRO002 | SQK-PCS111 | 151.04 | 970 |
| DTC_Patient 1 | ONT | No | N/A | FLO-PRO002 | SQK-PCS111 | 142.10 | 965 |
| DTC_Patient 1 | ONT | Yes | Immune Targeted+AM | FLO-PRO002 | SQK-PCS111 | 88.35 | 1250 |
| DTC_Patient 1 | ONT | Yes | Pan Cancer Targeted+AM | FLO-PRO002 | SQK-PCS111 | 87.3 | 1270 |
| DTC_Patient 2 | ONT | Yes | Immune Targeted+AM | FLO-PRO002 | SQK-PCS111 | 100.3 | 1300 |
| DTC_Patient 2 | ONT | Yes | Pan Cancer Targeted+AM | FLO-PRO002 | SQK-PCS111 | 95.26 | 1260 |

**Supplementary Table 2.** Differential transcript usage across dissociated tumor cell types.

| cell_type | cell_type | transcript_id | transcript_id | log2fc | log2fc | adj_pval | adj_pval |
| --- | --- | --- | --- | --- | --- | --- | --- |
| group1 | group2 | group1 | group2 | group1 | group2 | group1 | group2 |
| Epithelial | PDGFRa+/b+ Fibroblast | ENST00000551241_OAS1 | ENST00000202917_OAS1 | 25.14798355 | -8.292088509 | 1.539655352 | 303.8365371 |
| Epithelial | PDGFRa-/b+ Fibroblast | ENST00000433892_CD44 | ENST00000425428_CD44 | 28.92649651 | -5.610807419 | 19.19197313 | 17.14625666 |
| Epithelial | PDGFRa-/b+ Fibroblast | ENST00000579687_ICAM2 | ENST00000449662_ICAM2 | 26.62352371 | -6.31666851 | 4.272842283 | 20.23584951 |
| Epithelial | CD8+ Tcells | ENST00000559358_CD79B | ENST00000392795_CD79B | 26.9120388 | -5.32420969 | 5.253282315 | 19.17861045 |
| Epithelial | CD8+ Tcells | ENST00000587992_ICAM3 | ENST00000160262_ICAM3 | 5.071159363 | -7.313385487 | 37.90906839 | 119.9610031 |
| Epithelial | CD8+ Tcells | ENST00000397857_ITGB2 | ENST00000397846_ITGB2 | 26.7311039 | -5.332381248 | 5.065852457 | 104.8593928 |
| Epithelial | CD4+ Tcells | ENST00000397857_ITGB2 | ENST00000397846_ITGB2 | 26.7311039 | -5.534425735 | 5.06315725 | 121.5384365 |
| Epithelial | CD4+ Tcells | ENST00000317376_SPOCK2 | ENST00000536168_SPOCK2 | 26.99452972 | -6.621021748 | 5.269401875 | 172.6281979 |
| Epithelial | CD4+ Tcells | ENST00000513135_TNFRSF25 | ENST00000356876_TNFRSF25 | 25.02295876 | -5.524062634 | 1.421938819 | 74.24177565 |
| Epithelial | NK cells | ENST00000433892_CD44 | ENST00000425428_CD44 | 28.92649651 | -5.677530289 | 19.42820613 | 15.45279989 |
| Epithelial | NK cells | ENST00000559358_CD79B | ENST00000392795_CD79B | 26.9120388 | -5.064445972 | 5.305535154 | 14.89713269 |
| Epithelial | NK cells | ENST00000339438_CSDE1 | ENST00000533818_CSDE1 | 29.51158333 | -5.88548708 | 25.43342367 | 15.61530617 |
| Epithelial | NK cells | ENST00000558469_CUX1 | ENST00000547394_CUX1 | 26.16084099 | -5.287603855 | 3.13555046 | 15.15083082 |
| Epithelial | NK cells | ENST00000409817_CXCR4 | ENST00000241393_CXCR4 | 25.16540718 | -6.445358276 | 1.444525735 | 128.4669897 |
| Epithelial | NK cells | ENST00000589484_DAZAP1 | ENST00000590419_DAZAP1 | 26.48117447 | -5.757906914 | 4.214698382 | 15.52142723 |
| Epithelial | NK cells | ENST00000450253 EIF4E | ENST00000515638 EIF4E | 27.15531158 | -5.270740986 | 5.993981849 | 15.11424885 |
| Epithelial | NK cells | ENST00000610553_EWSR1 | ENST00000485037_EWSR1 | 28.91420555 | -5.237635612 | 18.72184672 | 15.10398688 |
| Epithelial | NK cells | ENST00000484194_HLA-E | ENST00000493699_HLA-E | 25.85902214 | -5.484964848 | 2.465774603 | 30.91073139 |
| Epithelial | NK cells | ENST00000587992_ICAM3 | ENST00000160262_ICAM3 | 30.55088043 | -7.022383213 | 44.04361269 | 106.5846861 |
| Epithelial | NK cells | ENST00000527146_IFITM2 | ENST00000399817_IFITM2 | 25.7280407 | -6.636305809 | 2.315677157 | 162.3786385 |
| Epithelial | NK cells | ENST00000534507_IL32 | ENST00000549213_IL32 | 29.8010788 | -6.055452347 | 29.0241439 | 31.97949804 |
| Epithelial | NK cells | ENST00000268182_IQGAP1 | ENST00000560218_IQGAP1 | 28.61159515 | -5.11726141 | 15.81584002 | 14.99118163 |
| Epithelial | NK cells | ENST00000482429_LTB | ENST00000429299_LTB | 26.13475037 | -5.074635029 | 3.152354308 | 108.4288332 |
| Epithelial | NK cells | ENST00000539616_MNAT1 | ENST00000553354_MNAT1 | 28.00305557 | -5.921472549 | 10.96362175 | 15.62535024 |
| Epithelial | NK cells | ENST00000478268_MX1 | ENST00000490220_MX1 | 28.45718193 | -5.452587605 | 14.10074914 | 15.26601429 |
| Epithelial | NK cells | ENST00000530902_PPP2R2B | ENST00000394409_PPP2R2B | 25.27556229 | -5.812587261 | 1.881190377 | 15.53450127 |
| Epithelial | NK cells | ENST00000537384_PRKAR1B | ENST00000400758_PRKAR1B | 27.86318207 | -5.326539516 | 9.722939295 | 45.87648056 |
| Epithelial | NK cells | ENST00000258062_REPS1 | ENST00000431346_REPS1 | 26.39013672 | -6.068675995 | 3.813781481 | 31.97346061 |
| Epithelial | NK cells | ENST00000536255_RPAIN | ENST00000574003_RPAIN | 33.34592438 | -5.759884834 | 127.0649907 | 15.48560643 |
| Epithelial | NK cells | ENST00000256196_RRAS2 | ENST00000531421_RRAS2 | 25.68914604 | -5.869750023 | 2.521183453 | 15.62973893 |
| Epithelial | NK cells | ENST00000478405_SPIN3 | ENST00000638619_SPIN3 | 29.61260796 | -5.892885685 | 27.20618712 | 15.62498037 |
| Epithelial | NK cells | ENST00000295754_TGFBR2 | ENST00000359013_TGFBR2 | 26.24829292 | -5.78046608 | 3.405474264 | 15.53227297 |
| Epithelial | DCs | ENST00000433892_CD44 | ENST00000425428_CD44 | 28.92649651 | -5.692490101 | 19.30584788 | 17.40599075 |
| Epithelial | DCs | ENST00000587992_ICAM3 | ENST00000160262_ICAM3 | 30.55088043 | -10.47030735 | 43.88204768 | 300.9619647 |
| Epithelial | DCs | ENST00000521636_IDO1 | ENST00000253513_IDO1 | 25.55454826 | -6.047535419 | 2.295939083 | 165.3045902 |
| Epithelial | DCs | ENST00000441671_LAG3 | ENST00000203629_LAG3 | 25.01807213 | -5.146058083 | 1.433436925 | 63.41837784 |
| PDGFRa+/b+ Fibroblast | CD8+ Tcells | ENST00000587992_ICAM3 | ENST00000160262_ICAM3 | 5.070431709 | -7.708373547 | 3.921013115 | 14.87540772 |
| PDGFRa+/b+ Fibroblast | CD4+ Tcells | ENST00000587992_ICAM3 | ENST00000160262_ICAM3 | 30.55015182 | -11.32203865 | 4.627280745 | 35.57702354 |
| PDGFRa+/b+ Fibroblast | CD4+ Tcells | ENST00000440815_IL32 | ENST00000533097_IL32 | 6.431498528 | -5.079892635 | 22.40176913 | 12.66753421 |
| PDGFRa+/b+ Fibroblast | NK cells | ENST00000587992_ICAM3 | ENST00000160262_ICAM3 | 30.55015182 | -7.417371273 | 4.688621162 | 13.1382252 |
| PDGFRa+/b+ Fibroblast | DCs | ENST00000587992_ICAM3 | ENST00000160262_ICAM3 | 30.55015182 | -10.86529541 | 4.598085166 | 37.96596885 |
| PDGFRa-/b+ Fibroblast | CD8+ Tcells | ENST00000440815_IL32 | ENST00000530890_IL32 | 5.615996838 | -5.253859997 | 8.621392831 | 5.940232191 |
| CD8+ Tcells | DCs | ENST00000372988_CCND3 | ENST00000372991_CCND3 | 30.15114212 | -5.235669613 | 3.021312019 | 10.23042604 |

**Supplementary Table 3.**
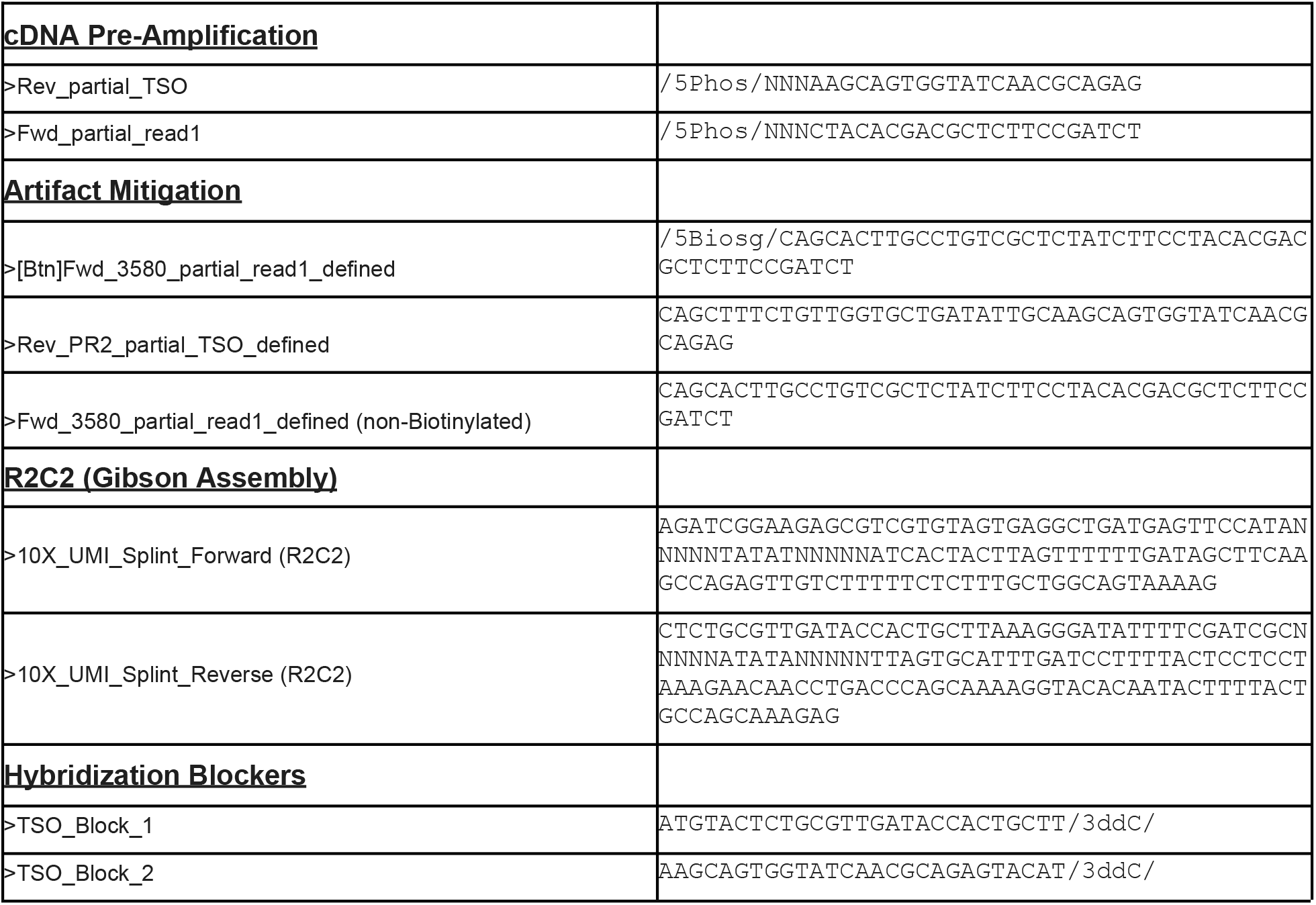
Primers and oligonucleotides used in this study. All oligonucleotides are shown in 5’ to 3’ direction and were ordered through Integrated DNA Technologies (IDT).

